# Activation of the mitochondrial unfolded protein response regulates the formation of stress granules

**DOI:** 10.1101/2023.10.26.564187

**Authors:** Marta Lopez-Nieto, Zhaozhi Sun, Emily Relton, Rahme Safakli, Brian D. Freibaum, J Paul Taylor, Alessia Ruggieri, Ioannis Smyrnias, Nicolas Locker

## Abstract

To rapidly adapt to harmful changes to their environment, cells activate the integrated stress response (ISR). This results in an adaptive transcriptional and translational rewiring, and the formation of biomolecular condensates named stress granules (SGs), to resolve stress. In addition to this first line of defence, the mitochondrial unfolded protein response (UPR^mt^) activates a specific transcriptional programme to maintain mitochondrial homeostasis. We present evidence that SGs and UPR^mt^ pathways are intertwined and communicate. UPR^mt^ induction results in eIF2α phosphorylation and the initial and transient formation of SGs, which subsequently disassemble. The induction of GADD34 during late UPR^mt^ protects cells from prolonged stress by impairing further assembly of SGs. Furthermore, mitochondrial functions and cellular survival are enhanced during UPR^mt^ activation when SGs are absent, suggesting that UPR^mt^-induced SGs have an adverse effect on mitochondrial homeostasis. These findings point to a novel crosstalk between SGs and the UPR^mt^ that may contribute to restoring mitochondrial functions under stressful conditions.

**Summary statement:** We describe a novel crosstalk between the mitochondrial unfolded protein response and the integrated stress response involving stress granules that protects cells from further stress.

## Introduction

Controlling the localisation and function of macromolecules is fundamental for many cellular functions and typically achieved by surrounding them with lipid membranes in organelles such as the nucleus, lysosomes, or mitochondria. In recent years, biomolecular condensates that lack membranes have become increasingly recognised as an alternative way to organise cellular components during stress. The dynamic formation of these condensates is supported by networks of protein-protein, protein- RNA and RNA-RNA interactions, generating high local concentration of RNA and protein components (Tauber et al., 2020). With such remarkable molecular plasticity, biomolecular condensates provide an ideal platform for the regulation of key processes, including mRNA metabolism or intracellular signalling, and are utilised by cells to rapidly adjust and rewire regulatory networks in response to various physiological and pathological triggers (Yoo et al., 2019).

Stress granules (SGs) are among the most characterised cytoplasmic biomolecular condensates and assemble in response to many stresses, including oxidative stress, heat shock, viral infection, proteasomal inhibition, endoplasmic reticulum (ER) stress and UV irradiation (Corbet and Parker, 2019; Hofmann et al., 2020). Capturing mRNAs and proteins, SGs are central to the reorganisation of cellular contents during stress. Stress sensing results in the activation of the integrated stress response (ISR) and a global inhibition of protein synthesis usually driven by phosphorylation of the eukaryotic translation initiation factor 2α (eIF2α), which results in the dissociation of mRNAs from polysomes and their accumulation in cytoplasmic RNP complexes (Hofmann et al., 2020). These RNPs are then recognised by aggregation prone RNA-binding proteins (RBPs), such as Ras-GTPase activating SH3 domain binding protein 1 (G3BP1) and T cell internal antigen-1 (TIA-1), which lead to further recruitment of multiple proteins containing low sequence complexity domains or intrinsically disordered regions. These mediate clustering and fusion events driven by multivalent interactions between their protein and RNA components, at the core of which G3BP1 acts as a key node that promotes RNA-protein, protein-protein and RNA-RNA interactions, ultimately resulting in liquid-liquid phase separation (LLPS) and SG formation (Corbet and Parker, 2019; Wang et al., 2018). SGs are highly dynamic, rapidly assembling to sequester the cytoplasmic stalled mRNAs and dissolving upon stress resolution to release stored mRNAs for future translation (Matheny et al., 2019; Namkoong et al., 2018). The dynamic nature of SGs is also reflected in their proposed stress-selective compositional, structural and functional heterogeneity, with different stresses proposed to induce at least 3 distinct types of SGs (Aulas et al., 2017; Hofmann et al., 2020). Type I and II SGs assemble via eIF2α-dependent and independent pathways, respectively, while type III SGs lack eIFs and are associated with cellular death (Reineke and Neilson, 2019). This suggests that compositionally heterogeneous SGs support specialized functions promoting survival or pro-death outcomes. By concentrating specific protein components and altering their cytoplasmic concentration, SGs have been proposed to alter the course of biochemical reactions and signalling cascades (Aulas et al., 2017; Hofmann et al., 2020). Furthermore, recent evidence hints that metabolic enzymes stored in SGs produce metabolites that can affect their stability (Begovich et al., 2020; Rollins et al., 2017; Saad et al., 2017). Importantly, dysregulation of pathways impacting SG clearance or LLPS is increasingly associated with neuropathology, in particular amyotrophic lateral sclerosis and related diseases (Wolozin and Ivanov, 2019). Moreover, SGs can be exploited by cancer cells to adapt to the tumour microenvironment, with resident SG proteins aberrantly expressed in tumours (Anderson et al., 2015). Finally, SGs are also proposed to be a crossroad between intracellular signalling and antiviral responses by controlling the location and function of key sensors and effectors of innate immunity (Eiermann et al., 2020).

In addition to this first line of defence, cells have evolved tailored quality control processes to protect the functions of key organelles in times of stress. Mitochondria act as the cell’s powerhouse, generating energy through oxidative phosphorylation but are also critical for essential function as such as calcium homeostasis, nucleotide metabolism and innate immunity regulation (Banoth and Cassel, 2018; Giorgi et al., 2018; Spinelli and Haigis, 2018). Key to maintaining or adapting mitochondrial functions, the mitochondrial unfolded protein response (UPR^mt^) is an evolutionarily conserved mitochondria-to-nucleus signalling cascade, initially described as a transcriptional response to the accumulation of misfolded mitochondrial proteins (Martinus et al., 1996; Zhao et al., 2002). Studies have shown that activation of the UPR^mt^ is primarily regulated by the Activating Transcription Factor 5 (ATF5) and Heat Shock Factor 1 (HSF-1) (Fiorese et al., 2016; Smyrnias et al., 2019; Sutandy et al., 2023; Wang et al., 2019), and engaged upon a wide range of mitochondrial insults, including abnormal level of mitochondrial reactive oxygen species (ROS) (Wang et al., 2022b), dissipation of mitochondrial membrane potential (Rolland et al., 2019), and reduced mitochondrial protein import efficiency (Nargund et al., 2012). Activated UPR^mt^ serves to restore mitochondrial protein homeostasis and promote cell survival in various biological processes, such as innate immune signalling (Pellegrino et al., 2014) and ageing (Houtkooper et al., 2013), as well as adaptation to pathophysiological conditions, such as neurodegenerative disorders (Sorrentino et al., 2017) and cardiovascular diseases (Smyrnias et al., 2019; Wang et al., 2019). Notably, growing evidence implicates the components of the ISR as central elements of the UPR^mt^ (Anderson and Haynes, 2020; Fu et al., 2023; Quirós et al., 2017).

The ISR is an adaptive gene expression regulation programme activated in response to a range of pathophysiological conditions, including hypoxia, amino acid deprivation, and viral infection (Brostrom et al., 1996; Dever et al., 1992; Harding et al., 2003; Ron, 2002; Wek et al., 2006). ISR signalling is initiated by the phosphorylation of eIF2α by four different kinases, which results in global translational shut down to dampen cellular workload and accumulation of SGs, while specific transcripts involved in stress adaptation remain preferentially translated. A key aspect of ISR regulation involves the fine-tuning of eIF2α phosphorylation by the growth arrest and DNA-damage inducible protein (GADD34). This stress-induced regulatory subunit of protein phosphatase 1 (PP1) mediates eIF2α dephosphorylation, acting as a negative feedback loop to prevent prolonged ISR activation to ensure stress resolution and resuming of protein synthesis (Harding et al., 2000; Lee et al., 2009; Novoa et al., 2001). Studying the components of the ISR signalling has provided evidence that the ISR acts as an essential precursor to the UPR^mt^ during mitochondrial stress. Specifically, each of the eIF2α kinases has been implicated in response to stress associated with mitochondrial dysfunctions (Baker et al., 2012; D’Souza and Minczuk, 2018; Guo et al., 2020; Samluk et al., 2019; Zhang et al., 2018). In addition, ATF5, together with ATF4 and C/EBP homologous protein (CHOP), are main regulators of the ISR that have emerged as essential mediators of a response to mitochondrial dysfunction similar to the UPR^mt^ in *C. elegans* (Fiorese et al., 2016; Jiang et al., 2020; Kaspar et al., 2021; Quirós et al., 2017; Zhao et al., 2002). Moreover, UPR^mt^ activation following mitochondrial dysfunction has been linked to reduced protein synthesis within the mitochondria (Münch and Harper, 2016), and with studies demonstrating that SGs impact metabolic processes, act as signalling hubs, and dock to mitochondria, we propose that SGs may be implicated in the interface between UPR^mt^ and ISR signalling during mitochondrial stress.

Herein, we established a novel crosstalk between the UPR^mt^ and SGs. Induction of UPR^mt^ during mitochondrial stress first results in an initial wave of SGs, which then disassemble potentially due to an upregulation of SG disassembly chaperones. Next, the negative feedback loop mediated by GADD34 protects cells against further stress insults and the assembly of eIF2α-dependent SGs. Finally, we demonstrated that the core SG scaffold proteins G3BP1 and G3BP2 (G3BP1/2) negatively affect mitochondrial respiration, morphology, ROS levels, and translation. Therefore, we propose a model in which the dynamic assembly of SGs is an important component of the UPR^mt^ to maintain mitochondrial homeostasis, which itself can fine-tune further adaptation to stress by activating GADD34. This provides further understanding of the cellular adaptation to adverse conditions and the crosstalk between generalised and specialised stress responses.

## Results

### UPR^mt^ activation by GTPP is associated with a transient SG assembly and translational shut-off

With previous studies suggesting an interplay between the UPR^mt^ and the ISR, and given the central role of SGs in the cellular stress response, we investigated the impact of mitochondrial stressors on the SG pathway in more details. First, we tested the induction of UPR^mt^ signalling in U2OS cells following mitochondrial stress using the mitochondrial HSP90 inhibitor gamitrinib-triphenylphosphonium (GTPP), which results in protein misfolding in the mitochondria and induction of mitochondrial chaperones and proteases (Kang et al., 2009). The levels of typical transcripts induced upon UPR^mt^ and ISR activation were measured using RT-qPCR (Fig 1A). We observed a clear induction of CHOP and ATF4 after 4 h of GTPP treatment, which are related to ISR activation, as well as HSPD1 and LONP1, indicating induction of the UPR^mt^. ATF5 acts as a key mediator of the mammalian UPR^mt^ signalling, translocating to the nucleus upon stress to activate a specific transcriptional cascade (Fiorese et al., 2016). Consistently, siRNA-mediated silencing of ATF5 expression significantly reduced the transcription of all GTPP-induced UPR^mt^ markers, but not ISR markers, further supporting the role of ATF5 as UPR^mt^ inducer in our model (Fig 1A).

**Figure 1:**
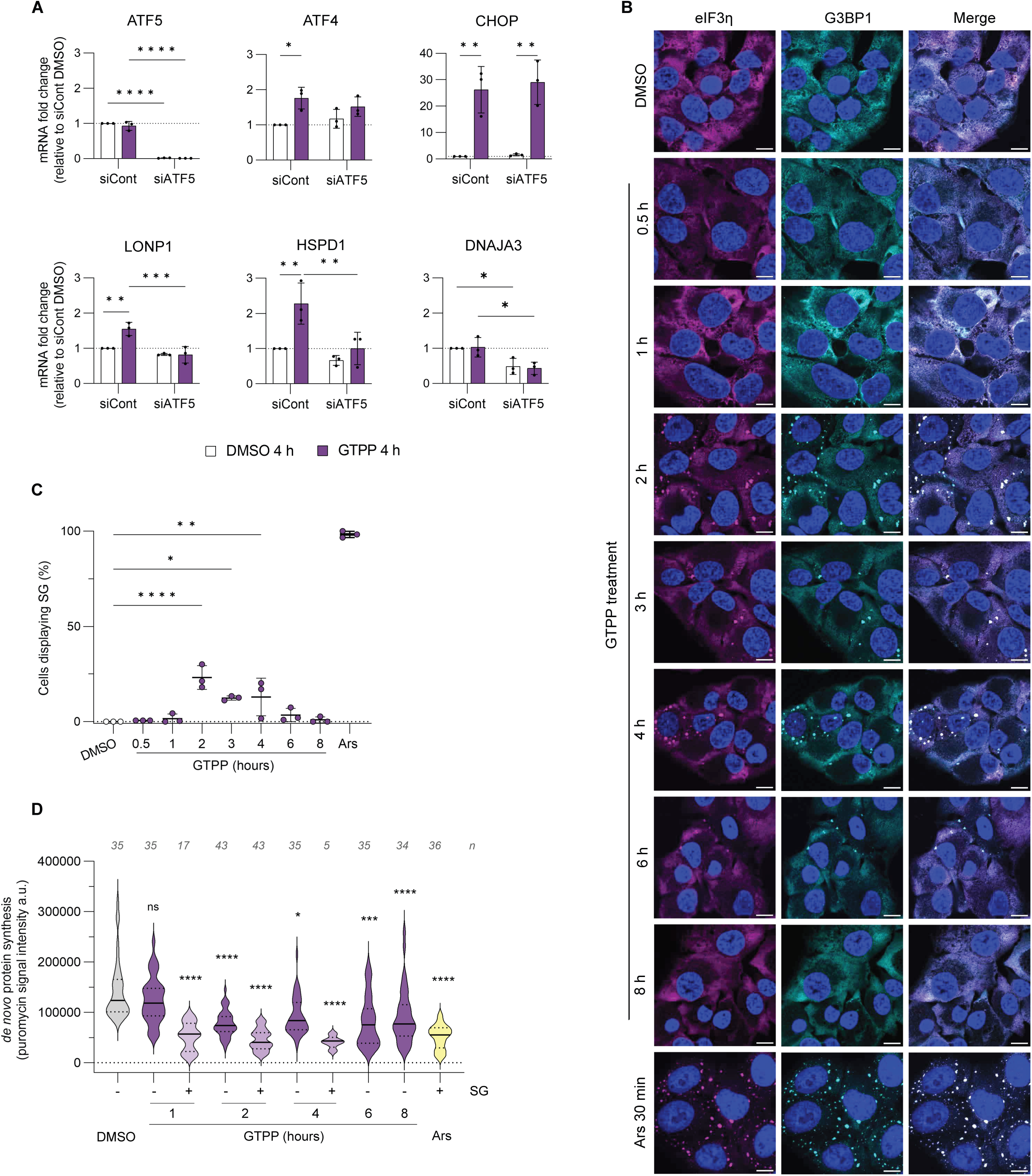
ATF5-dependent UPR^mt^ induction by GTPP is associated with early SG formation and translational shut off. (**A**) U2OS cells were transfected with non-targeting siRNA (siCont) or ATF5-specifc siRNA (siATF5). After 48 h cells were treated either with DMSO or 10 μM GTPP for 4h and mRNA levels of UPR^mt^ markers were measured by RT-qPCR. Results shown as mean ± SD, n=3, normalised to RPL9 mRNA and shown relative to siCont DMSO expression level, analysed by two-way ANOVA (****, P < 0.0001; ***, P < 0.001; **, P < 0.01; *, P < 0.05). (**B**) Cells were treated with GTPP up to 8 h and analysed by immunofluorescence for the SG markers G3BP1 (cyan) and eIF3η (magenta). Nuclei were stained with DAPI. Scale bars indicate 10 μm. (**C**) Quantification by manual counting of cells with SGs using ImageJ, at least 100 cells were counted per replicate. Results are shown as mean ± SD, n=3 and were analysed by one-way ANOVA compared to DMSO treated cells (****, P < 0.0001; *, P < 0.05). (**D**) Representative violin plots from three independent experiments showing *de novo* protein synthesis upon GTPP treatment over time. Arsenite (Ars)-treated cells were included as positive control, shown in yellow. Cells positive for SGs are shown in light purple. Figure shows the median and quartiles of puromycin signal intensity measured by immunofluorescence, number of cells analysed from this replicate is shown above per each condition and were analysed by one-way ANOVA compared to DMSO treated cells (****, P < 0.0001; **, P < 0.01; *, P < 0.05). Representative images are shown in Sup Fig 2.

Next, we tested the impact of UPR^mt^ activation on SG assembly by treating U2OS cells with GTPP for up to 8 hours. Cells were fixed at the indicated time points and stained with anti-G3BP1 and anti-eIF3η antibodies to detect SG assembly (Fig 1B). Sodium arsenite was used as positive control for SG assembly as it induces eIF2α-dependent SGs by activating the eIF2α kinase HRI (Taniuchi et al., 2016). G3BP1 and eIF3η remained diffused in the cytoplasm in DMSO-treated cells and, as expected, arsenite treatment resulted in the assembly of SGs reflected by the accumulation of G3BP1/eIF3η into cytoplasmic foci. Interestingly, approximately 25% of cells were positive for SGs at 2 h post GTPP treatment (Fig 1C). The percentage of SG-positive cells decreased gradually until being undetectable at 8 h post treatment. To further explore the characteristics of GTPP-induced SGs, we used recently described small molecules inhibitors of G3BP1 condensation that impair SG assembly and induce their disassembly (Freibaum et al., 2024). Pre-treatment of cells with the active inhibitor G3Ib completely blocked GTPP-induced SG formation, while the inactive compound G3Ib’ had no impact. This confirmed that SGs induced following UPR^mt^ activation, like canonical SGs, are dependent on G3BP1 condensation for their assembly (Sup Fig 1). Thus, mitochondrial stress following GTPP treatment in U2OS cells activates the UPR^mt^ and induces an early and transient phase of SG assembly.

Several studies have suggested a connection between mitochondrial function and cytosolic protein translation (Lu and Guo, 2020; Topf et al., 2018). Given that formation of SGs is preceded by an inhibition of translation, we examined the impact of GTPP on global translational control at a single cell level using ribopuromycylation assays (David et al., 2012). Briefly, cells were treated with arsenite as positive control or GTPP for the indicated time and nascent polypeptides were labelled with puromycin, a tRNA analogue, followed by an emetine pulse to avoid the release of puromycin-nascent chains from ribosomes. Cells were then fixed and stained with anti-puromycin to detect puromycylated nascent peptide chains, and with the SG markers G3BP1 and eIF3η (Sup Fig 2). Overall, GTPP treatment resulted at in strong translation repression at 2, 4, 6 and 8h post treatment when compared to untreated cells (Fig 1D). In addition, in treated cells displaying SGs, translational repression was more potent than in cells not forming SGs (Fig 1D). Therefore, these results indicate that UPR^mt^ activation by GTPP significantly attenuates cytosolic translation at 2 h post treatment and this shut off is at least in partial uncoupled from SG formation and maintained over time.

### GTPP-mediated mitochondrial stress leads to ISR activation by PERK

Phosphorylation of eIF2α is central event in the ISR signalling and a trigger of SG formation (Kedersha et al., 2002). Thus, we measured phosphorylation levels of eIF2α as well as protein and RNA expression levels of ISR markers during early UPR^mt^ activation. Phosphorylation of eIF2α was detected within 30 minutes of GTPP treatment, which was followed by an increase of ATF4 and GADD34 protein levels at 1 h and 3 h post treatment (Fig 2A). Similarly, CHOP and GADD34 mRNA levels were also highly upregulated following GTPP treatment, for 2 and 3 h, respectively (Fig 2B). These results indicate that UPR^mt^ induction by GTPP signals through an early activation of canonical ISR.

**Figure 2:**
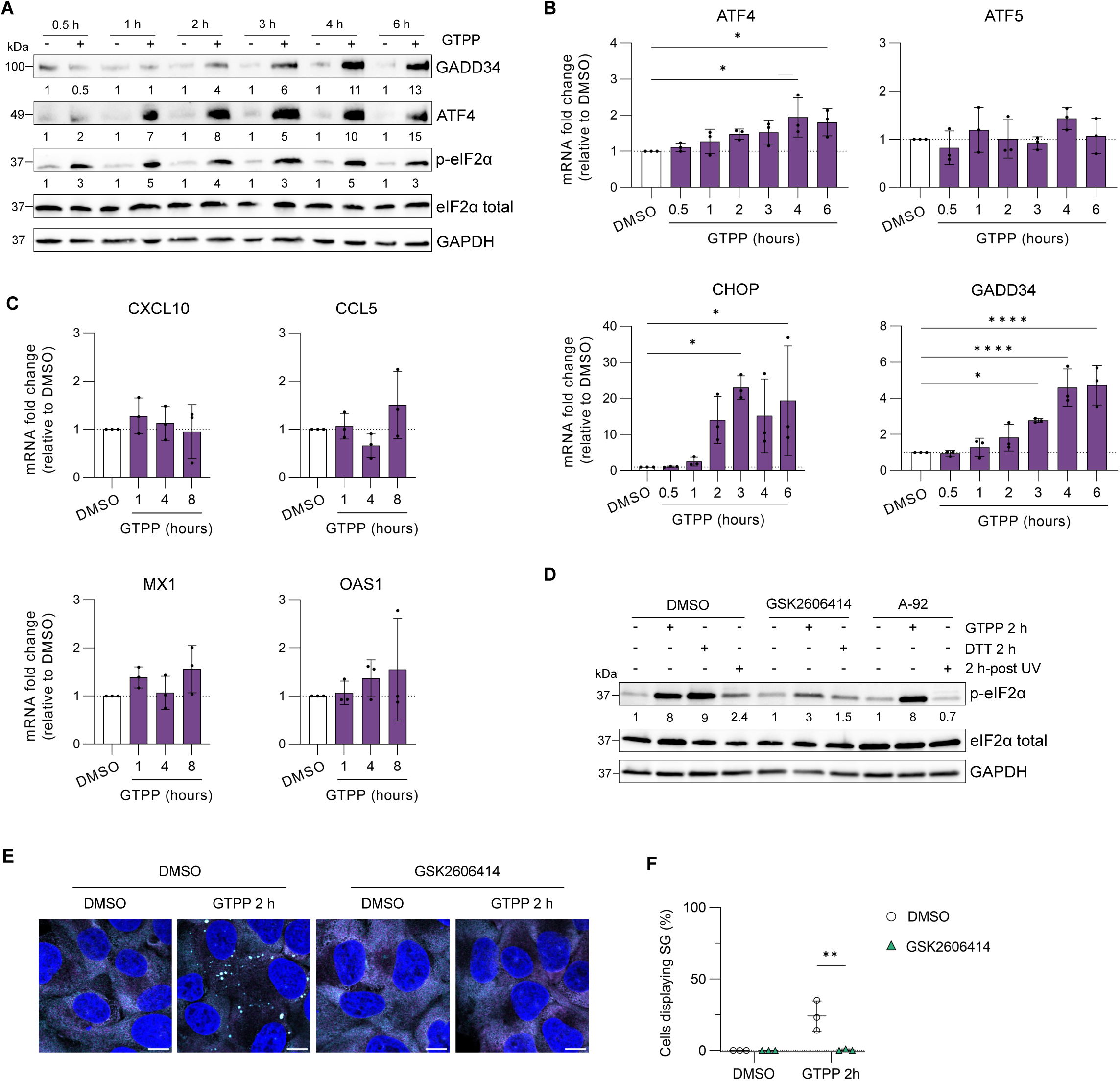
UPR^mt^ activation by GTPP results in activation of the integrated stress response (ISR) through PERK. (**A**) Representative western blot of three independent experiments showing protein levels of GADD34, ATF4, and p-eIF2α in U2OS cells treated with GTPP for up to 6. Band intensities were normalised to that of GAPDH and to DMSO for each time point. (**B**) RT-qPCR analysis of ATF4, ATF5, CHOP and GADD34 mRNA levels in U2OS cells treated with GTPP for up to 6 h. Results shown as mean ± SD, n=3, normalised to RPL9 mRNA and shown relative to DMSO expression level, analysed by one-way ANOVA (****, P < 0.0001; *, P < 0.05). (**C**) RT-qPCR analysis of the interferon-stimulated genes CXCL10, CCL5, MX1 and OAS1 after 1, 4 and 8 h GTPP exposure. Results shown as mean ± SD, n=3, normalised to RPL9 mRNA and shown relative to DMSO expression level. (**D**) Representative immunoblot from three independent experiments showing levels of p-eIF2α in U2OS cells treated with either DMSO, 0.1 μM PERK inhibitor GSK2606414 or 1 μM GCN2 inhibitor A-92 for 1 h, followed by treatment with 10 μM GTPP for 2 h, or 5 mM DTT for 2 h, or 2 h-post UV light exposure at 20mJ/cm^3^. Band quantifications were normalised to GAPDH and relative to DMSO for each inhibitor used. (**E**) Cells were pre-incubated with either DMSO or 0.1 μM PERK inhibitor GSK2606414, further stimulated with GTPP for 2 h and analysed by immunofluorescence for the SG markers G3BP1 (cyan) and eIF3η (magenta). Nuclei were stained with DAPI. Scale bars indicate 10 μm. (**F**) Quantification by manual counting of cells with SGs using ImageJ, at least 100 cells were counted per replicate. Results are shown as mean ± SD, n=3 and were analysed by two-way ANOVA compared to DMSO treated cells (**, P < 0.001).

We next tested which of the four eIF2α kinases is activated during UPR^mt^ induction. We first hypothesized that the double-stranded RNA (dsRNA)-activated protein kinase (PKR) might be activated upon GTPP treatment due to mitochondrial dsRNA leakage occurring as a consequence of mitochondrial stress and membrane damage (Kim et al., 2018). However, mitochondrial dsRNA was not detected after UPR^mt^ induction (Sup Fig 3). Supporting this result, the expression of the interferon-stimulated genes such as CCXCL10, CCL5, MX1 and OAS1, reported to be induced upon PKR activation (Pindel and Sadler, 2011; Radetskyy et al., 2018), was not upregulated during early or late UPR^mt^ (Fig 2C). Thus, PKR is unlikely to be the kinase responsible for eIF2α phosphorylation upon GTPP treatment. We next investigated the potential role of PKR-like ER kinase (PERK) and general control non-derepressible protein 2 (GCN2). PERK is activated in response to ER stress and has been shown to modulate mitochondrial homeostasis during stress (Almeida et al., 2022), while GCN2 is activated by amino acid depravation or UV light radiation (Deng et al., 2002). We used the pharmacological inhibitors GSK2606414 and A-92 to inhibit PERK and GCN2 activity in U2OS cells, respectively. We first confirmed that they reduced eIF2α phosphorylation upon DTT treatment or UV light exposure (Fig 2D). In response to 2 h GTPP, the GCN2 inhibitor A-92 did not reduce eIF2α phosphorylation, whereas the PERK inhibitor GSK2606414 decreased it by more than 2-fold, suggesting that PERK is the main eIF2α kinase responsive to GTPP. To confirm that PERK activation is responsible for SG assembly during UPR^mt^ activation, cells were treated with PERK inhibitor followed by GTPP treatment. This resulted in the complete lack of SGs further confirming that GTPP- induced SGs are dependent on PERK-driven eIF2α phosphorylation (Fig 2E, 2F). Overall, these results suggest that activation of the UPR^mt^ results is rapidly followed by an early phase characterised by ISR activation through PERK and SG assembly (0.5-3 h), and a late phase where SGs disassemble and UPR^mt^ genes are expressed (4-8 h).

### Mitochondrial stress impairs the assembly of eIF2α-dependent SGs

As GTPP-mediated mitochondrial stress and UPR^mt^ activation resulted in SG assembly only during the early and not at the later stages of stress, we hypothesized that the UPR^mt^ may limit the ability of cells to form SGs during prolonged stress. To test this, cells were stimulated for 2, 4, 6 or 8 h with GTPP and then challenged with arsenite for 30 min to induce the formation of SGs (Fig 3A and Sup Fig 4). As expected, in absence of pre-treatment with GTPP, arsenite induced the assembly of SGs in almost all cells. Interestingly, only some GTPP-treated cells formed SGs after arsenite treatment at the indicated timepoints. Detailed analysis of U2OS cells revealed that pre-treatment with GTPP reduced arsenite-induced SG assembly compared to arsenite-only stimulated cells from 92% to 50%, 14%, 28% and 35% after 2, 4, 6 or 8 h treatments, respectively (Fig 3B). These data indicate that UPR^mt^ induction by GTPP inhibits subsequent arsenite-induced SG formation. SG assembly is a multistep process by which microscopically invisible “seeds” first coalesce upon stress, and then fuse into larger aggregates during a growth phase (Wheeler et al., 2016). Thus, we next examined whether the SG maturation process is also affected when UPR^mt^ is pre-activated. To address this, the average size of SGs per cell and number of SGs per cell were compared in cells pre-treated with GTPP and non-treated (Fig 3C). The analysis showed that while a 2 h GTPP treatment does not impact the area of arsenite-induced SGs, longer pre-treatment results in much smaller ones significantly smaller (in average 0.47, 0.53, 0.5 μm^2^, for 4 h, 6 h and 8 h respectively) compared to non-treated cells (in average 0.72 μm^2^). Likewise, less arsenite-induced SGs were detectable in GTPP pre-treated cells. These results suggest that SG maturation process and fusion into larger granules may be altered by GTPP during late UPR^mt^.

**Figure 3:**
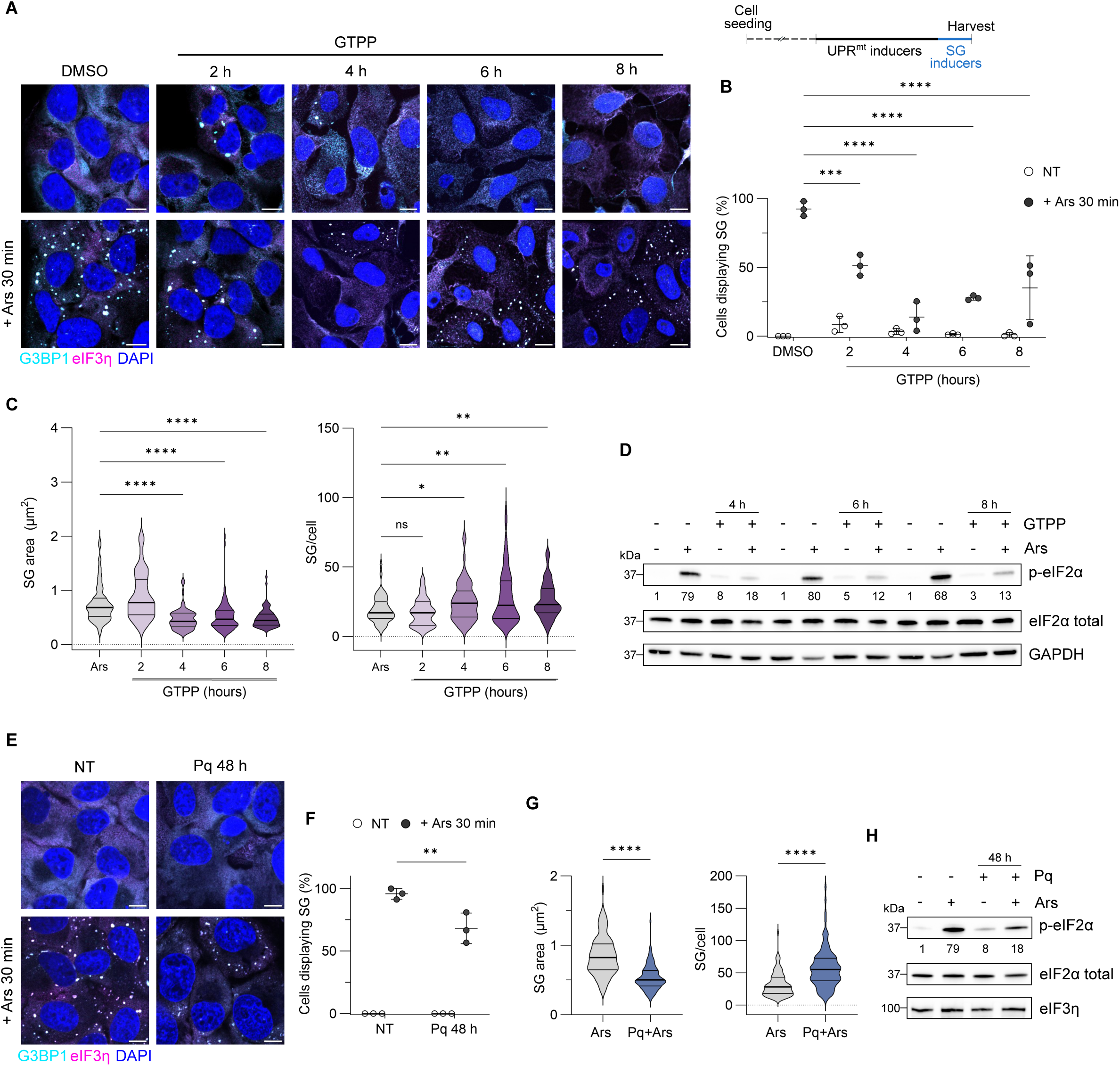
Pre-treatment with UPR^mt^ inducers impairs assembly of eIF2α-dependent SGs and prevents eIF2α phosphorylation. Representative immunofluorescence images (**A, E**) and quantified data (**B, F**) of U2OS cells pre-treated with 10 μM GTPP **(A, E**) or 0.5 Paraquat **(B, F)**. Sodium arsenite (0.5 mM) was added 30 min before harvesting of cells to induce SG formation (schematic diagram of the experimental setup is shown on the top right). Cells were analysed by immunofluorescence for the SG markers G3BP1 (cyan) and eIF3η (magenta). Nuclei were stained with DAPI. Scale bars indicate 10 μm. Quantification was done by manual counting of cells with SGs using ImageJ and at least 100 cells were counted per replicate. Results are shown as mean ± SD, n=3 and were analysed by two-way ANOVA (****, P < 0.0001; ***, P < 0.001). The average size of SGs per cell and the number of SGs per cell in U2OS cells treated with GTPP (C) or Paraquat (G) were quantified using the G3BP1 channel and Analyze Particle plugin from ImageJ,, as displayed as violin plot displaying showing the median and quartiles of at least 60 cells per condition from three independent experiments analysed by (C) Krustal-Wallis test (*, P < 0.05; **, P < 0.01; (****, P < 0.0001) and (G) by Mann-Whitney test (****, P < 0.0001). (**D, H**) Representative blot from three independent experiments showing p-eIF2α protein levels in U2OS cells treated as in (A)and (E).. eIF2α phosphorylation levels were normalized to GAPDH expression levels and are shown as relative to DMSO/NT cells; NT, non-treated.

Given that eIF2α phosphorylation drives SG formation, we next assessed how UPR^mt^ activation impacts on arsenite-induced eIF2α phosphorylation. Cells were treated as mentioned above and levels of basal and phosphorylated eIF2α measured by immunoblotting. Cells treated with arsenite displayed high levels of eIF2α phosphorylation, while GTPP treatment slightly increased eIF2α phosphorylation as shown previously (Fig 3D). Interestingly, when subjected to arsenite treatment, GTPP pre-treated cells showed lower levels of arsenite-induced eIF2α phosphorylation. Specifically, phosphorylated levels were reduced approximately 3 to 4-fold compared to arsenite-treated cells in all timepoints.

To confirm these results, UPR^mt^ was induced with another mitochondrial stressor, paraquat, which produces high amounts of superoxide and interferes with the electron transport chain (Cocheme and Murphy, 2008). Treatment of cells with paraquat for 48 h increased by 2 to 3-fold the mRNA levels of ATF4, CHOP, GADD34 and DNAJA3 compared to non-treated cells, confirming ISR and UPR^mt^ activation (Sup Fig 5A). Next, we examined how paraquat stimulation impacts on SG assembly and whether pre-treatment with paraquat affects the ability of cells to form arsenite-induced SGs by immunofluorescence. Paraquat treatment for up to 48 h did not induce SG formation, and G3BP1 or eIF3η remain diffused in the cytoplasm (Sup Fig 5B). However, prior induction of UPR^mt^ by treatment with paraquat for 48 h decreased percentage of cells containing arsenite-induced SGs, from about 96% to 65% (Fig 3E, 3F and Sup Fig 5C). Similar to our previous observation with GTPP, paraquat pre-treatment resulted in smaller and less numerous arsenite-induced SGs when compared to non-treated (0.53 μm^2^ compared to 0.85 μm^2^, and 57 vs 32, respectively) (Fig 3G). These results indicate that paraquat impairs SG formation and limits their maturation into larger SGs. Paraquat treatment also resulted in a slight increase in eIF2α phosphorylated levels, supporting ISR induction, and pre-treatment with paraquat also limited the levels of arsenite-induced eIF2α phosphorylation compared to non-treated cells (Fig 3H). Overall, activation of UPR^mt^ using two different stressors results in the inhibition of arsenite-induced eIF2α phosphorylation and consequently prevents SG formation during prolonged stress. Altogether, these data strongly propose that SG assembly is modulated by prior induction of the UPR^mt^.

Given that ATF5 is one of the central regulators in UPR^mt^ activation, we explored its impact on SG assembly (Fiorese et al., 2016). To study this, cells were transfected with siRNA targeting ATF5 and SG formation was monitored by immunofluorescence during GTPP treatment for 4 h and with or without arsenite (Sup Fig 6A). ATF5 silencing did not affect SG assembly nor disassembly after 4 h of GTPP treatment, around 20% of cells were positive for SGs (Sup Fig 6B). Moreover, assembly of arsenite-induced SGs in GTPP pre-treated cells was impaired when ATF5 was silenced, with no differences compared to non-silencing control cells. However, quantitative analyses revealed that GTPP-induced SGs were significantly smaller when ATF5 was silenced compared to control cells, while no difference was observed in the number of granules per cell (Sup Fig 6C). In contrast, whereas ATF5 silencing did not altered the size of arsenite-induced SGs, it resulted in a significant reduced number of granules compared to non-silencing cells (Sup Fig 6D). SGs differ in their dynamics, assembly and composition depending on the stress (Aulas et al., 2017), therefore these findings suggest that the ATF5-mediated UPR^mt^ transcriptional reprogramming alters SG dynamics in a stress-specific manner.

### GADD34 is required for the inhibition of *de novo* eIF2α-dependent SG assembly by preventing eIF2α phosphorylation

We observed that UPR^mt^ activation by either GTPP or paraquat results in the upregulation of GADD34 (Fig 2A, 2B and Sup Fig 5A), the negative regulator of the ISR, which in complex with PP1 dephosphorylates eIF2α to resume translation. Therefore, we hypothesized that increased levels of GADD34 could limit eIF2α phosphorylation in response to a consecutive stress and consequently SG assembly. To test this hypothesis, we generated U2OS GADD34 KO cell clones using a CRISPR/Cas9 approach (Sup Fig 7A). Control cell clones (Ctrl) were generated for comparison using a non-targeting crRNA. Ctrl and GADD34 KO cells were stimulated with GTPP for 4 h prior arsenite treatment and SG assembly was detected by immunofluorescence (Fig 4A and Sup Fig 7B). As observed previously, prior induction of UPR^mt^ by GTPP impaired the assembly of arsenite-induced SGs in Ctrl cells. In contrast, we observed no difference in the assembly of SGs in GADD34 KO cells irrespectively of GTPP pre-treatment. While only 17% of Ctrl cells pre-treated with GTPP were positive for arsenite-induced SG, 97% of GADD34 KO cells contained SGs in the same conditions (Fig 4B). Similar results were obtained with paraquat pre-treatment with 66% Ctrl cells and 98% GADD34 KO cells displaying SGs in response to arsenite treatment (Fig 4C, 4D and Sup Fig 7C).

**Figure 4:**
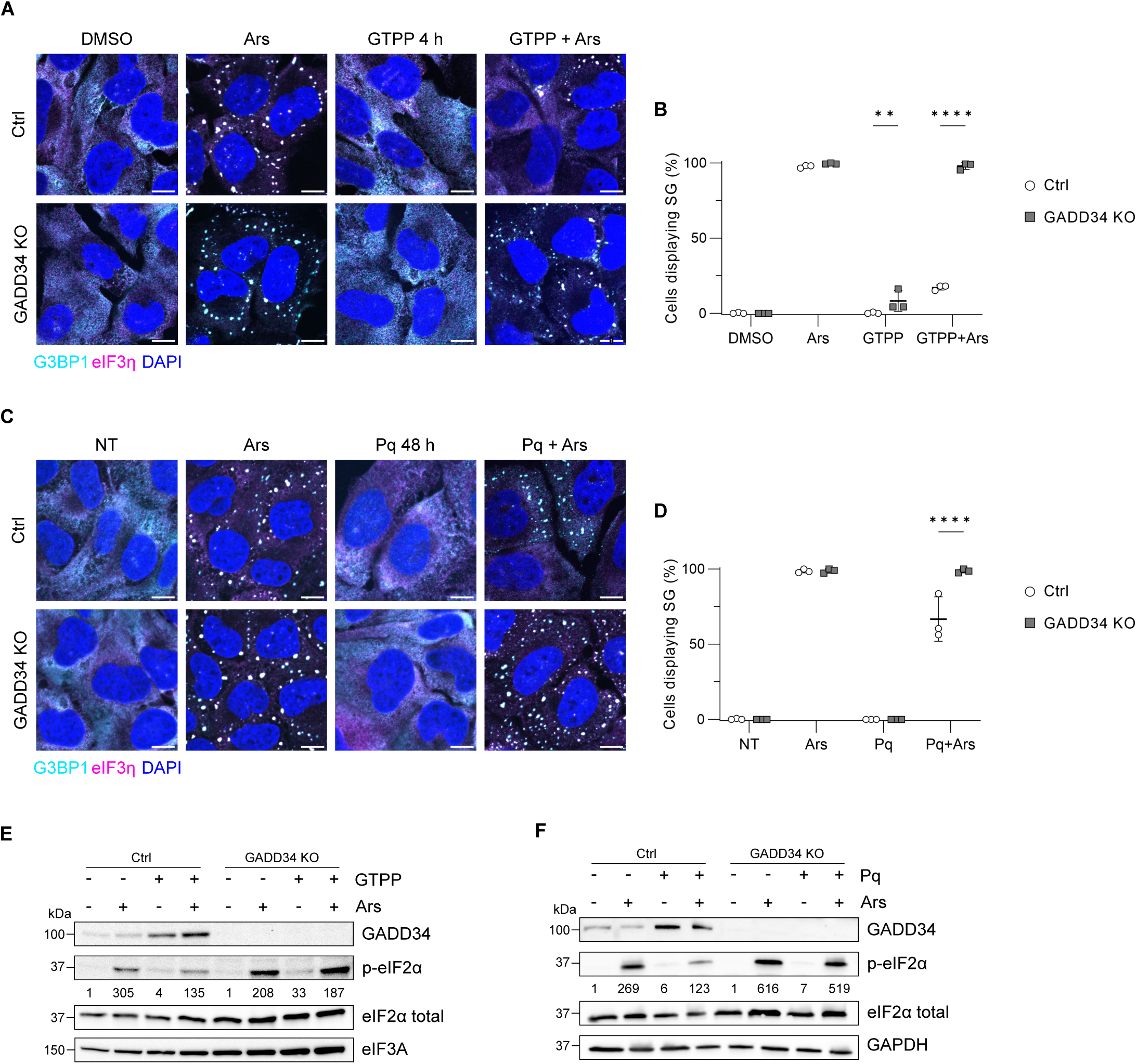
GADD34 upregulation is responsible for preventing eIF2α-phosphorylation and SG assembly. U2OS control (Ctrl) or GADD34 KO cells were treated with GTPP 4 h (**A**) or paraquat (Pq) 48 h (**C**) and 0.5 mM sodium arsenite (Ars) was added 30 min before harvest and cells were analysed by immunofluorescence for the SG markers G3BP1 (cyan) and eIF3η (magenta). Nuclei were stained with DAPI. Scale bars indicate 10 μm. (**B, D**) Panels show quantification by manual counting of cells with SGs using ImageJ, at least 100 cells were counted per replicate. Results are shown as mean ± SD, n=3 and were analysed by two-way ANOVA (****, P < 0.0001; **, P < 0.01). (**E, F**) Representative western blot analysis of three independent experiments of same conditions as above showing GADD34 protein levels and arsenite-induced eIF2α phosphorylation levels in Ctrl and GADD34 U2OS cells pre-treated with GTPP (**E**) and Pq (**F**). eIF2α phosphorylation levels were normalized to eIF3A or GAPDH expression levels and are shown as relative to DMSO/NT cells; NT, non-treated.

These results were supported by reduced levels of phosphorylated eIF2α in GTPP and paraquat pre-treated Ctrl cells compared to GADD34 KO cells (Fig 4E, 4F). In Ctrl cells, either GTPP or paraquat treatment followed by arsenite stimulation resulted in reduced eIF2α phosphorylation levels compared to stimulation with arsenite only. In contrast, in GADD34 KO cells, the induction of eIF2α phosphorylation was comparable between stimulation with arsenite and GTPP or paraquat treatments followed by arsenite stimulation. Therefore, GADD34 KO rescued the ability of cells to phosphorylate eIF2α following UPR^mt^ induction. Overall, these results indicate that UPR^mt^ activation results in increased levels of GADD34 expression, which prevent eIF2α phosphorylation and thereby impair the ability of cells to assemble SGs during stress adaptation.

### GTPP-mediated UPR^mt^ activation differentially modulates the assembly of eIF2α-independent SGs

SG assembly can also be triggered following eIF2α-independent inhibition of translation, for example by blocking eIF4A or mTOR activity (Aulas et al., 2017). Thus, we examined whether UPR^mt^ activation alters the assembly of eIF2α-independent SGs following treatment with the eIF4A inhibitor silvestrol (Bordeleau et al., 2008). To test this, U2OS cells were stimulated with GTPP for 6 h, then treated with or without silvestrol for 1 h and SG assembly was analysed by immunofluorescence (Fig 5A). As expected, silvestrol induced formation of SGs in nearly all cells. Interestingly, pre-treatment with GTPP resulted in a reduction of the percentage of cells displaying silvestrol-induced SGs, from 94% to 72% (Fig 5B). In contrast, pre-treatment with paraquat did not impair the assembly of silvestrol-induced SGs. (Fig 5D, 5E). However, features of SGs appeared to be altered by both mitochondrial stressors, as reported before for arsenite-induced SGs. Detailed analysis revealed that GTPP and paraquat treatment resulted in a reduction in average size of silvestrol-induced SGs, from 0.98 μm^2^ to 0.9 μm^2^ and 0.66 μm^2^, respectively, and increased average number of SGs, from 18 to 22 and 39, respectively, when compared to non-treated cells (Fig 5C, 5F). These results indicate that UPR^mt^ activation by GTPP impairs the ability of cells to assemble eIF2α-independent SGs, although not to the same extent as eIF2α-dependent SGs, and that both stressors affect their maturation and fusion process. Taken together, these data suggest that UPR^mt^ activation impact on SG assembly and dynamics both in an ISR-dependent and -independent manner.

**Figure 5:**
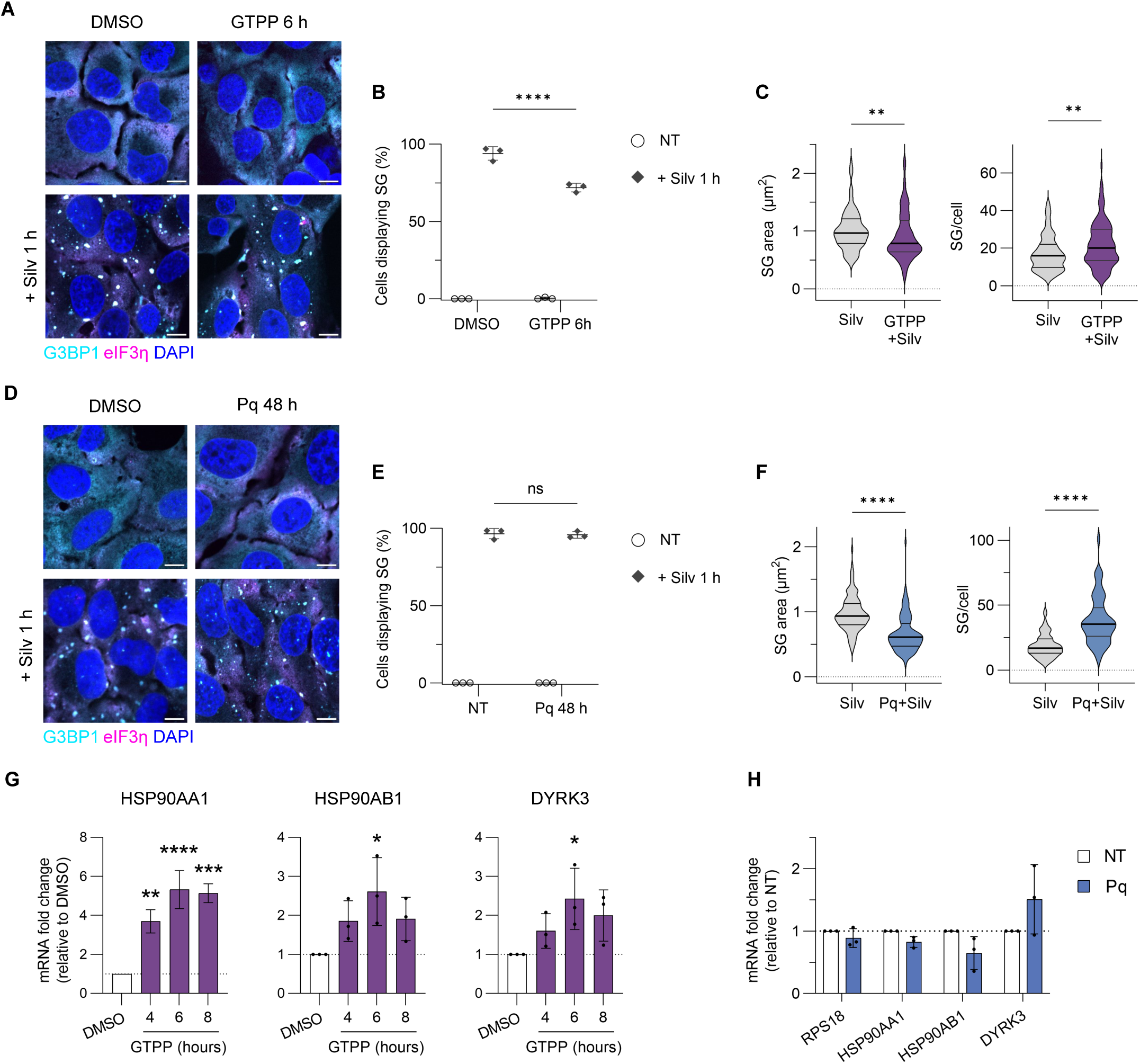
UPR^mt^ activation also alters eIF2α-independent SG formation associated with an upregulation of SG disassembly chaperones. U2OS were pre-treated with GTPP (**A**) or paraquat (Pq) (**D**) and 2 μM silvestrol (Silv) was added 1 h before harvest, analysed by immunofluorescence for the SG markers G3BP1 (cyan) and eIF3η (magenta). Nuclei were stained with DAPI. Scale bars indicate 10 μm. (**B, E**) Panels show quantification by manual counting of cells with SG using ImageJ, at least 100 cells were counted per replicate. Results are shown as mean ± SD, n=3, and were analysed by two-way ANOVA (****, P < 0.0001; ns, non-significant). (**C, F**) Violin plot displaying the average size of SGs per cell and the number of SGs per cell of data from A and D, respectively, using G3BP1 channel and Analyze Particle plugin from ImageJ. Figures show the median and quartiles of at least 100 cells per condition from three independent experiments and were analysed by Mann-Whitney test (****, P < 0.0001; **, P < 0.01). (**G, H**) RT-qPCR analysis of SG disassembly chaperones HSP90AA1, HSP90AB1 and DYRK3 mRNA expression levels in response to GTPP (**G**) and Pq (**H**) treatment. Results shown as mean ± SD, n=3, normalised to RpL9 mRNA and shown relative to DMSO/NT expression level, analysed by two-way ANOVA (****, P < 0.0001; **, P < 0.01; *, P < 0.05).

SGs are highly dynamic and disassemble upon stress removal. Recently, the chaperone Hsp90 was reported to initiate SG disassembly by interacting with and activating the dual specificity tyrosine-phosphorylation-regulated kinase 3 (DYRK3), leading to the phosphorylation of yet unknown targets. (Mediani et al., 2021). Given that GTPP-induced SGs disassemble after 4 h treatment (Fig 1C) and to further examine how UPR^mt^ activation modulates assembly and dynamics of eIF2α-independent SGs, we measured the mRNA levels of DYRK3 and the stress-inducible and constitutively expressed Hsp90 forms, HSP90AA1 and HSP90AB1, respectively. Upon UPR^mt^ induction by GTPP, the level of HSP90AA1 mRNA was significantly increased by approximately 4 to 6-fold in all time points compared to non-treated cells, while Hsp90AB1 and DYRK3 were slightly upregulated by 2 to 3-fold at 6 and 8 h GTPP treatment (Fig 5G). In contrast, the mRNA levels of these proteins remained unchanged upon UPR^mt^ induction by paraquat (Fig 5H). These results are consistent with our previous results where silvestrol-induced SG assembly was altered by GTPP treatment but not by paraquat, indicating that GTPP but not paraquat mediated UPR^mt^ activation enhances SG disassembly process. Altogether, these data suggest that UPR^mt^ activation by GTPP results in an initial SG assembly which then rapidly disassemble potentially due to an enhancement in the SG disassembly process.

### Cells unable to assemble SGs during stress present improved mitochondrial functions during UPR^mt^ <u>activation</u>

To investigate the impact of reduced SG formation during mitochondrial stress, we first examined the effect of GTPP on mitochondrial morphology in U2OS G3BP1/2 double knockout cells (G3BP1/2 dKO) lacking the ability to form SGs compared to control cells (Sup Fig 8) (Yang et al., 2020). Wild-type (WT) and G3BP1/2 dKO U2OS cells were treated with GTPP for 2 h, the time at which we previously detected the appearance of SGs, and stained for the mitochondrial marker TOMM20 (Fig 6A). Confocal imaging of DMSO-treated cells showed a typical tubular mitochondrial network while those treated with GTPP contained round and fragmented mitochondria. Subsequent detailed analysis indicated that following GTPP treatment, 91.5% of WT cells presented fragmented mitochondria, compared to 75% of G3BP1/2 dKO cells (P < 0.012) (Fig 6B). These results indicate that GTPP-induced defective mitochondrial dynamics is partially restored when the SG scaffolding proteins G3BP1/2 are absent. Alterations in mitochondrial morphology are associated with mitochondrial ROS (mtROS) and in particular, high mtROS levels promote mitochondria fragmentation or swelling (Brillo et al., 2021). We then studied whether GTPP results in mtROS production and how the absence of SGs impacts on mtROS levels. WT and G3B1/2 dKO cells were stimulated with GTPP for 2 h and incubated with MitoSOX fluorogenic dye which specifically detects mitochondrial superoxide in live cells. Interestingly, while GTPP treatment induced high production of mtROS in WT cells, 1.7-fold compared to non-treated, GTPP-induced mtROS levels were significantly lower in G3BP1/2 dKO cells (Fig 6C and Sup Fig 9). These data demonstrate that loss of G3BP1/2 partially prevents mitochondria fragmentation and reduces mtROS production during mitochondrial stress.

**Figure 6:**
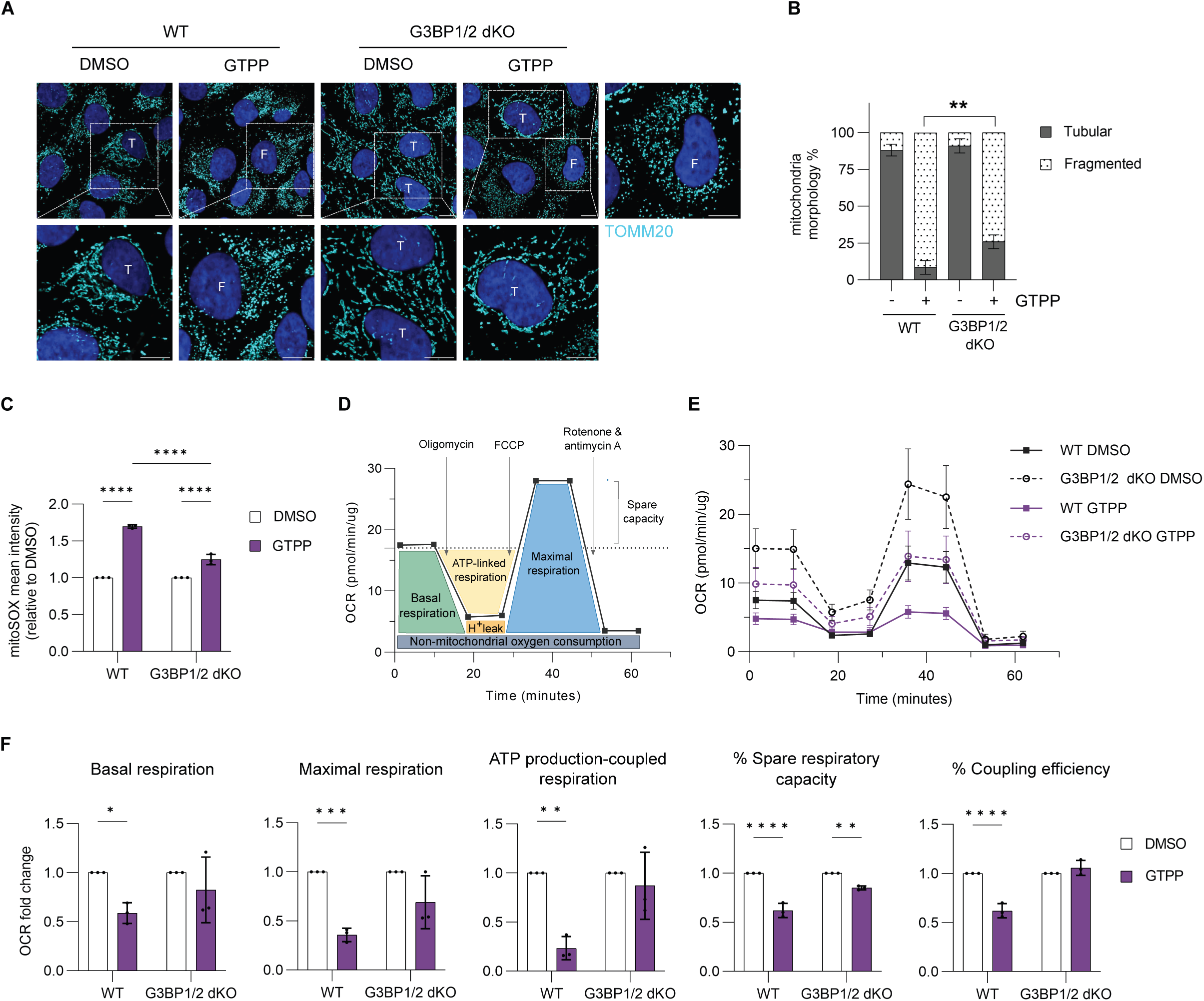
Presence of SGs alters mitochondrial functions during UPR^mt^. (**A**) WT and G3BP1/2 dKO cells were treated with either DMSO or GTPP for 2 h and analysed by immunofluorence for the mitochondrial marker TOMM20 (cyan). Nuclei were stained with DAPI. Scale bars indicate 10 μm. T: tubular mitochondria; F: fragmented mitochondria. (**B**) Quantification of the mitochondrial phenotypes observed in **A**, at least 100 cells were counted per biological replicate. Figure shows mean ± SD, n=3 and analysed by two-way ANOVA (**, P < 0.01). (**C**) Following GTPP treatment for 2 h, U2OS WT and G3BP1/2 cells were incubated with 2.5 uM MitoSOX dye for 30 mins, and mitochondrial ROS were quantitatively measured using IncuCyte S3 live cell analysis software. Figure shows mean ± SD, n=3, relative to DMSO-treated cells, and analysed by two-way ANOVA (****, P < 0.0001). (**D**) Schematic diagram of the respiratory profile of live cells using Seahorse Cell Mito Stress Test, showing the key parameters of mitochondrial function and the drugs being used. Briefly, oxygen consumption rate is measured when cells are sequentially stimulated with oligomycin (ATP synthase inhibitor), FCCP (protonophore, uncoupling) and antimycin A/rotenone (inhibitors of complex III and I). (**E-F**) Representative traces (**E**) and averaged OCR data (**F**) from three independent experiments in WT (solid line) or G3BP1/2 dKO (dashed line) U2OS cells treated with DMSO (black) or GTPP (purple) for 2 h. Results shown as mean ± SD, n=3 and as fold change to DMSO-treated cells for each cell line, analysed by two-way ANOVA (****, P < 0.0001; ***, P < 0.001; **, P < 0.01; *, P < 0.05).

To further characterise the functional consequences of reduced SG formation during mitochondrial stress, we compared mitochondrial respiration upon UPR^mt^ activation in WT and G3BP1/2 dKO cells. Cells were treated with GTPP for 2 h and measured mitochondrial oxygen consumption rate (OCR) using a Seahorse flux analyser (Fig 6D). As expected GTPP impaired mitochondrial functions in WT cells resulting in decreased basal, maximal respiration, ATP production-coupled respiration and spare respiratory capacity compared to untreated cells (Fig 6E, 6F). In contrast, GTPP-stressed G3BP1/2 dKO cells that cannot form SGs presented with significantly improved mitochondrial respiration when compared to WT cells.

Protein synthesis is regulated during stress conditions to preserve cellular resources. Given that GTPP stimulation results in arrest of cytosolic translation (Fig 1D), we next investigated how GTPP impacts on translational control when G3BP1/2 are absent. G3BP1/2 dKO cells were treated with GTPP, or arsenite as a control, for the indicated times and ribopuromycylation assay was performed as described above to measure the levels of translation in single cells. As shown in previous studies (Ohn et al., 2008), arsenite treatment induced a potent translational shut off both in WT and G3BP1/2 dKO cells, confirming that SG formation is not required for translation repression (Fig 7A, 7B). Interestingly, in G3BP1/2 dKO cells GTPP treatment did not alter the levels of translation which contrasts with WT cells (Fig 1D and Fig 7A, 7B), suggesting that the translational shut off induced by GTPP is dependent on the presence of G3BP1/2 and/or the formation of UPR^mt^-induced SGs. Next, we hypothesise that the impact of GTPP treatment on protein synthesis could be linked to specific translational control at the mitochondrial level. To further assess this, we monitored the impact of GTPP treatment, on specific mitochondrial translation using a SUNSET-based approach (Yousefi et al., 2021). Cells were incubated in methionine-free media and cytosolic translation was selectively halted by cycloheximide treatment, before adding the alkyne-containing non-canonical amino acid L-homopropargylglycine (HPG). This allows HPG to be specifically incorporated into mitochondrial translation products, instead of methionine, which can be visualized by copper-catalyzed cycloaddition reaction (click) to azide-containing fluorescent dyes (Fig 7C, 7D, 7E). First, we confirmed the validity of this approach as the HPG-derived fluorescence signal colocalise with mitochondrial TOMM20 signal. Second, when cells were left in methionine-containing media or incubated with the mitochondrial translation inhibitor chloramphenicol (CHL), HPG signal disappeared showing that HPG is only incorporated into mitochondrial nascent peptides, and its fluorescence signal correlates with mitochondrial translation (Fig 7C). Quantification of HPG incorporation revealed that UPR^mt^ activation by GTPP results in decreased mitochondrial translation in WT cells. Interestingly, G2BP1/2 dKO cells also presented a significant reduction in mitochondrial translation after stimulation with GTPP (Fig 7D, 7E), indicating that absence of G3BP1/2 does not affect mitochondrial translation during UPR^mt^ as compared to cytosolic translation. Taken together, these results demonstrated that the assembly of UPR^mt^-induced SGs is associated with altered mitochondrial dynamics, mtROS production without impairing mitochondrial translation during stress.

**Figure 7:**
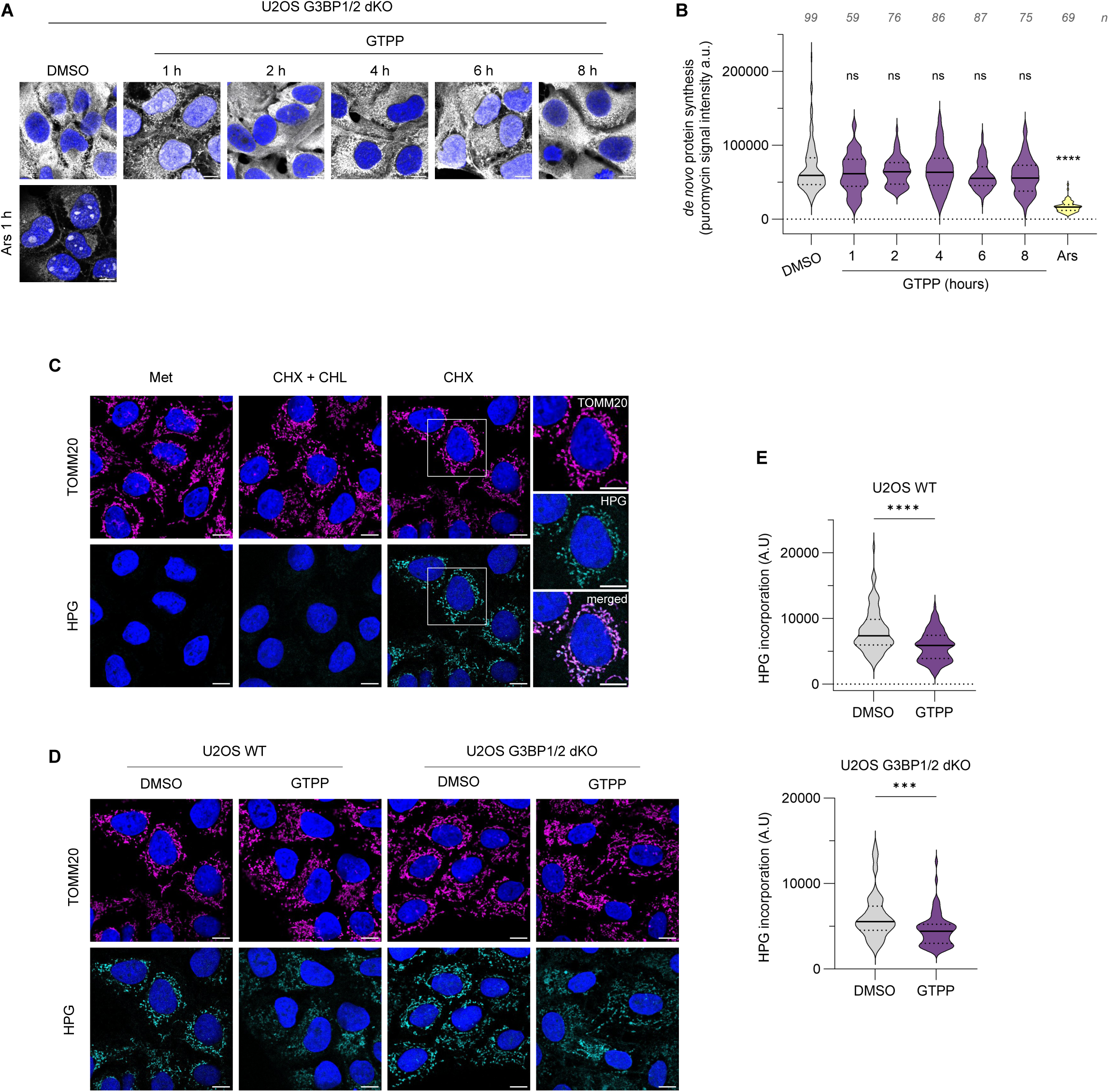
G3BP1/2 are required for effective cytosolic translational shut off but not mitochondrial during UPR^mt^ activation. (**A**) G3BP1/2 dKO cells were treated with GTPP or arsenite for the indicated times and stimulated with puromycin followed by emetine before fixation. Then, cells were analysed by immunofluorescence with anti-puromycin (grey). Nuclei were stained with DAPI. Scale bars indicate 10 μm. (**B**) Representative violin plots from three independent experiments showing *de novo* protein synthesis upon GTPP treatment over time. Arsenite (Ars)-treated cells were included as positive control, shown in yellow. Figure shows the median and quartiles of puromycin signal intensity measured by immunofluorescence, number of cells analysed from this replicate is shown above per each condition and were analysed by one-way ANOVA compared to DMSO treated cells (****, P < 0.0001). (**C**) U2OS WT cells were either incubated in normal media (Met, methionine) as a control, or methionine-free media in the presence and/or absence of cycloheximide (CHX) and/or chloramphenicol (CHL), before adding HPG for 15 minutes. After fixation, AF555-azide (cyan) was used for click-chemistry reaction to visualise HPG incorporation and cells were additionally labelled with TOMM20 as mitochondrial marker (magenta). Nuclei stained with DAPI. Scale bars indicate 10 μm. (**D**) WT and G3BP1/2 dKO cells were treated with either DMSO or GTPP for 2h, before inhibiting cytosolic translation with cycloheximide and performing HPG labelling. After fixation, cells were fixed, labelled and imaged as in (**C**). Scale bars indicate 10 μm. (**E**) Representative violin plots from three independent experiments showing HPG incorporation upon 2h GTPP treatment. For quantification, images were subjected to thresholding before AF555 signal intensity was measured using measure function in ImageJ. Results were analysed by unpaired t-test for each cell line compared to DMSO treated cells (***, P < 0.001; ****, P < 0.0001).

### Loss of SG scaffolding proteins G3BP1/2 results in enhanced cell survival during chronic UPR^mt^ <u>activation</u>

Mitochondrial homeostasis is crucial for cell survival and prolonged mitochondrial dysfunction can lead to cell death (Van Houten et al., 2016). We therefore examined how the absence of SGs impacts on cell survival following chronic UPR^mt^ activation. The kinetics of cell viability after GTPP treatment in WT and G3BP1/2 dKO U2OS cells was monitored for 28 h using the IncuCyte live-cell imaging system. Interestingly, we detected increased resistance to GTPP-induced cell death in G3BP1/2 dKO cells that are unable to form SGs, when compared to WT cells (Fig 8). The number of WT dead cells, indicated by round-shape orange-fluorescent cells, increased significantly compared with WT DMSO treated cells at 11 h of GTPP treatment (Fig 8A, 8B). In contrast, we did not detect a significant increase in G3BP1/2 dKO dead cells until 17 h after GTPP treatment compared to DMSO-treated cells. These results show that cells unable to form SGs present a prolonged cell death following UPR^mt^ activation compared to WT cells, suggesting that GTPP-induced SGs are detrimental for cellular survival.

**Figure 8:**
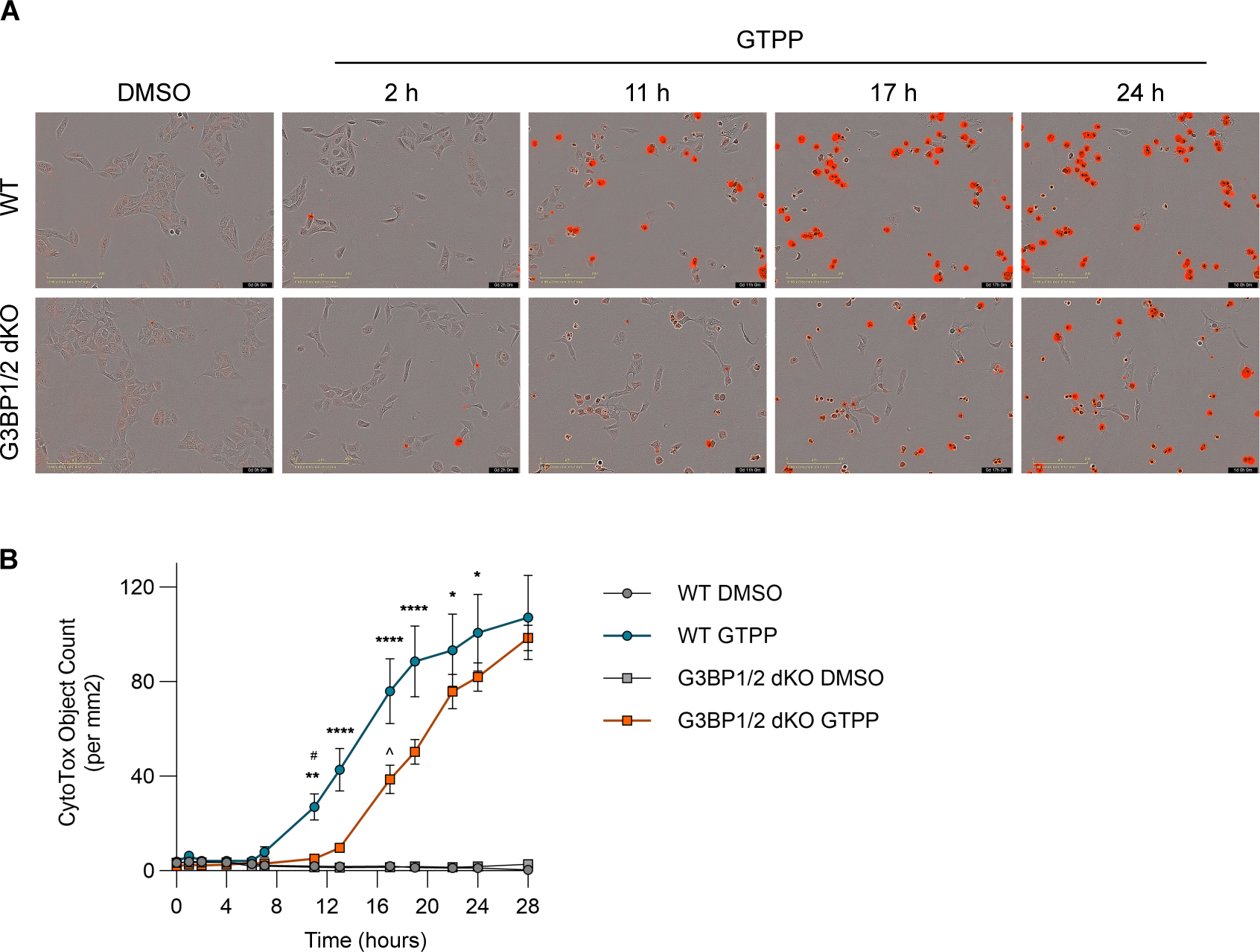
Loss of SG scaffolding proteins G3BP1/2 delays cell death following prolonged UPR^mt^ activation. GTPP and Cytotox Dye were added to WT and G3BP1/2 U2OS cells and monitored for 28 h using IncuCyte S3 live cell analysis software. (**A**) Selected representative time frames of DMSO and GTPP-treated cells. Scale bar indicates 200 μm. (**B**) Representative cell viability kinetics of three independent experiments of WT and G3BP1/2 dKO cells when exposed to GTPP. The figure shows mean ± SEM of cell death count (orange cytotox dye) from 9 images per condition. Data were analysed by two-way ANOVA (****, P < 0.0001; **, P < 0.01; *, P < 0.05; when comparing WT GTPP vs G3BP1/2 dKO GTPP. #, P < 0,001; when comparing WT DMSO vs WT GTPP. ^, P < 0.0001; when comparing G3BP1/2 dKO DMSO vs G3BP1/2 dKO GTPP).

## Discussion

The disruption of mitochondrial homeostasis is associated with several pathologies, such as neurodegeneration, cancer, aging, and cardiovascular diseases (Chen et al., 2023; Norat et al., 2020; Quirós et al., 2016; Suomalainen and Battersby, 2018). The UPR^mt^ is an evolutionarily conserved homeostatic pathway engaged upon a wide range of mitochondrial insults, including mitochondrial protein misfolding (Zhao et al., 2002), reduced mitochondrial protein import efficiency (Nargund et al., 2012), dissipation of mitochondrial membrane potential (Donehower et al., 2019), and dysregulated levels of reactive oxygen species (ROS) (Wang et al., 2022b). UPR^mt^ activation triggers a broad and protective transcriptional cascade, through enhancing mitochondrial folding capacity and triggering a mitochondrial recovery program, alleviating neurodegenerative and cardiovascular diseases symptomatology (Smyrnias et al., 2019; Sorrentino et al., 2017), promoting resistance to pathogen infection (Mahmud et al., 2020; Pellegrino et al., 2014), and prolonging lifespan (Munkácsy and Rea, 2014). Studies in mammals have shown that the ISR and UPR^mt^ are intertwined; however, how SGs – which are downstream effectors of the ISR – modulate the outcome of the UPR^mt^ and how their assembly is regulated upon mitochondrial stress is unknown. Evidence supporting a functional interplay between SGs and mitochondria include the impaired metabolism linked to altered SG dynamics in several neurodegenerative diseases, the regulation of SG stability by metabolites produced by resident SG enzymes such as Sam1 or pyruvate kinase, and the regulation of mitochondrial permeability and lipid fatty acid oxidation by starvation-induced SGs (Begovich et al., 2020; Cereghetti et al., 2021; Rollins et al., 2017; Saad et al., 2017; Wang et al., 2022a). SGs can physically contact different membrane-bound organelles, including lysosomes or the ER, and starvation-induced SGs contact with mitochondria in human cells (Amen and Kaganovich, 2021; Lee et al., 2020; Liao et al., 2019). Furthermore, numerous mitochondrial proteins have been associated with SGs (Amen and Kaganovich, 2021; Aoyama-Ishiwatari et al., 2021; Paget et al., 2023). Our study identifies SGs as novel signalling components associated with the mammalian UPR^mt^ and SG dynamics as a key for maintaining mitochondrial function during mitochondrial stress. We show that UPR^mt^ activation by GTPP or paraquat results in PERK-mediated activation of the ISR, accompanied with transient SG formation, in the case of GTPP. Moreover, UPR^mt^-induced upregulation of the GADD34 protects cells against persistent SG assembly and preventing SG formation improved mitochondrial functions upon UPR^mt^ induction.

ATF5 has been proposed to drive the protective effect of UPR^mt^ in different diseases models such as osteoarthritis (Zhou et al., 2022), heart failure (Smyrnias et al., 2019; Wang et al., 2019), and protect host during enteric infection (Chamseddine et al., 2022). In our experimental system, we show that ATF5 is essential for the upregulation of the UPR^mt^ markers in cells subjected to mitochondrial stress (Fig 1), but while it was not involved directly in SG formation, GTPP-induced SGs were smaller when ATF5 was silenced (Sup Fig 6). Thus, impaired upregulation of UPR^mt^ genes may affect the composition and maturation of SGs, as well as their function. Recently, a study by Sutandy *et al*. proposed that the accumulation of mitochondrial protein precursors and ROS in the cytosol results in nuclear translocation of heat shock factor 1 (HSF1) and activation of UPR^mt^ markers. Importantly, they demonstrated that ATF5 synergises with HSF1, which affects late activation of the UPR^mt^ (Sutandy et al., 2023). Thus, we cannot exclude the possibility that other transcription factors such as HSF1 may contribute to SG dynamics during conditions of mitochondrial stress and induction of the UPR^mt^.

The disassembly of SGs is crucial for cellular homeostasis since persistent SGs can promote the formation of aberrant aggregates, which are implicated in several neurodegenerative diseases (Baron et al., 2013; Repici et al., 2019; Vanderweyde et al., 2012). SGs induced by GTPP begin to disassemble post 3 h treatment and completely disappear by 8 h (Fig 1C). We first hypothesized that the increased expression of GADD34 due to ISR activation might be responsible for SG disassembly in these mid/late phases. However, GADD34 KO cells cleared SGs with a comparable kinetic to control cells (Fig 5), indicating that a factor other than GADD34 is involved in SG disassembly after GTPP treatment. In addition, GTPP treatment resulted in increased expression of the kinase DYRK3, and HSP90, a major chaperone promoting SG disassembly of SGs through binding and stabilising DYRK3 (Fig 5) (Mediani et al., 2021; Wippich et al., 2013). Upregulation of SG disassembly chaperones was not detected with paraquat, consistent with this stressor not inducing SG formation. Interestingly, UPR^mt^ induction by either GTPP or paraquat prevented *de novo* assembly of arsenite-induced SGs by impairing eIF2α phosphorylation. This effect was rescued in GADD34 KO cells, indicating that GADD34 upregulation is responsible for the inhibition of SG assembly following a second stressor (Fig 4). This result is in agreement with recent work demonstrating that sustained GADD34 levels upon ISR activation reduces cell responsiveness to a second stress pulse, promoting cell adaptation (Klein et al., 2022; Magg et al., 2024). Our data suggest that UPR^mt^ employs GADD34 as a negative regulator of the ISR to fine-tune cellular stress responses and maintain mitochondrial function under these conditions.

UPR^mt^ activation by GTPP also inhibited the formation of eIF2α-independent SGs, and, the size and number of both eIF2α-dependent and independent SGs were altered in cells treated with both GTPP and paraquat compared to non-treated. This suggests that UPR^mt^-induced stress could alter SG biogenesis by limiting the growth of small SGs prior to their fusion into bigger aggregates. We speculate that the known GTPP and paraquat effects on oxidative phosphorylation could affect SG biogenesis, whose fusion and growth steps are ATP-dependent (Jain et al., 2016). Further experiments are needed to fully understand how energy depletion affects SG biogenesis during mitochondrial stress.

Our results reveal that cells subjected to mitochondrial stress activate the UPR^mt^ and engage mechanisms to disassemble either SGs induced by mitochondrial stress and/or prevent *de novo* assembly of SGs, suggesting that clearance of SGs is crucial to the maintenance of mitochondrial homeostasis. Consistently, we show that GTPP-induced SGs impaired mitochondrial morphology and respiration, whereas cells unable to form SGs exhibited improved mitochondrial function, including translation during stress (Fig 6). In addition, absence of SGs improved cell survival during mitochondrial stress (Fig 8). Furthermore, GTPP treatment blocked cytosolic translation uncoupled from SG formation and when G3BP1/2 were absent, this translational shut off was rescued (Fig 7). Besides eIF2α phosphorylation, protein synthesis is also modulated by the mammalian target of rapamycin (mTOR) signalling pathway. Activation of mTOR leads to hyperphosphorylation of 4E-BP which allows the correct assembly of eIF4F complex into the 5’-cap and thereby translation initiation. Insufficient nutrients or energy result in mTOR inhibition, preventing cap-dependent translation (Haghighat et al., 1995; Zoncu et al., 2011). UPR^mt^ activation by GTPP reduces ATP production-coupled respiration (Fig 6F) which may inhibit mTOR resulting in translation inhibition independent of SG formation. G3BP1/2 have been described as anchors of the tuberous sclerosis complex (TSC) to lysosomes and supress mTOR activity upon starvation, thus, G3BP1 silencing results in mTOR hyperactivation during those conditions (Prentzell et al., 2021; Rehbein et al., 2021). Therefore, an hyperactivation of mTOR signalling pathway in G3BP1/2 dKO cells could explain the unaltered translation levels after UPR^mt^ activation. Further studies are required to unravel the molecular mechanisms underlying translational control during UPR^mt^ activation and disentangle the effect driven by SG assembly from those of G3BP1 itself.

Questions remain as to the functional link between inhibition of SG assembly and mitochondrial homeostasis following UPR^mt^ activation. Several studies support a protective role for SGs, through impairing pro-apoptotic JNK signalling (Arimoto et al., 2008; Kitajima et al., 2023), sequestration of caspases (Fujikawa et al., 2023) or suppressing production of ROS (Park and Shin, 2023; Takahashi et al., 2013) and they have been described as “shock absorbers” dampening excessive innate immune response to viral infection (Paget et al., 2023). In contrast, persistent SGs are implicated in pro-death functions during chronic stress conditions as well as neurodegenerative diseases or tumour chemotherapy (Reineke et al., 2018; Reineke and Neilson, 2019; Schwed-Gross et al., 2022). Future studies are required to further compositionally characterise SGs, and their functions, during mitochondrial stress.

Overall, our findings support a model in which mitochondrial stress-induced SGs are formed as a consequence of the associated ISR activation. Subsequently, cells activate homeostatic mechanisms to disassemble SGs*, i.e.* the UPR^mt^, as their persistence may compromise mitochondrial recovery. This could occur by through various mechanisms, including physically preventing efficient mitochondrial dynamics, recruiting mitochondrial or related proteins to SGs, preventing translation of essential mitochondrial proteins, or a combination of these. This underscores a novel functional interplay between the ISR and UPR^mt^ and further work should aim at revealing the impact of SG signalling during UPR^mt^, how UPR^mt^-induced SGs, and the sequestration of specific resident proteins, impact on mitochondrial homeostasis. This will open further opportunities for alternative strategies for the treatment of human diseases with compromised mitochondrial function and associated with an impaired SG dynamics.

## Material and methods

### Cell culture and gene silencing

U2OS, U2OS G3BP1/2 double knock out (dKO), U2OS control (Ctrl) and GADD34 KO cell lines were grown in Dulbecco’s modified Eagle’s medium (DMEM) supplemented with 10% fetal bovine serum, 100 U/ml penicillin, 100 g/ml streptomycin and 1% L-glutamine in a humidified incubator at 37°C and 5% CO_2_ environment. U2OS G3BP1/2 dKO cells have been described previously (Burke et al., 2020).

Silencing of ATF5 was performed using Lipofectamine-RNAiMax transfection reagent (Thermo Fisher). U2OS cells were incubated with the transfection mix containing either siRNA negative control (Invitrogen, #4390843) or siRNA targeting ATF5 (Invitrogen, ID s22425) at a final concentration of 10 nM for 48 h. Next, cells were incubated with subsequent treatments and analysed as described below.

### Generation of U2OS GADD34 KO cell clones

U2OS GADD34 KO cells were generated using a two-crRNA strategy to delete a portion of the *PPP1R15A* locus encoding GADD34. crRNAs cutting sequences in *PPP1R15A* exon 2 (IDT, Hs.Cas9.PPP1R15A.AC: AltR1 / AUC ACU GGG UUG GCA CCA CCG UUU UAG AGC UAU GCU /AltR2) and exon 3 (IDT, custom design: AltR1 / GAG GCG UGG CUG AGA CCA ACG UUU UAG AGC UAU GCU / AltR2) were purchased to Integrated DNA technologies (IDT). Individual crRNAs and tracrRNA (IDT) were resuspended in 1 x TE buffer (10 mM Tris, 0.1 mM EDTA, pH 7.5) to a final concentration of 200 µM and hybridized at equimolar amounts by heating to 95 °C for 5 min and slow stepwise cooling (1°C/sec) to room temperature. Pre-assembled crRNA/Cas9 ribonucleoprotein complexes (RNPs) were obtained by gently mixing 1.2 µl of hybridized RNAs with 17 µg of recombinant Alt-R Cas9 protein (IDT) and 2.1 µl of PBS, followed by 20 min incubation at room temperature.

U2OS cells (1 x 10^6^) were washed with PBS, resuspended in 91 µl nucleofection solution from Cell Line Nucleofector Kit V (Lonza) and mixed with 2.5 µl of each pre-assembled RNPs and 4 µl electroporation enhancer (Integrated DNA technologies). Nucleofection was performed using an Amaxa 2b nucleofector (Lonza) and pre-set program X-01. Nucleofected cells were incubated for 48 h prior to clonal amplification and screening for homozygous clones using target-specific PCR (GoTaq Hot Start, Promega) on genomic DNA using the following primers: PPP1R15A_forward: 5’-ATCACACCGGGAGTGTTGTC-3’, PPP1R15A_reverse: 5’-AGACGCTGCTCGCTACAAAT-3’) flanking sequences targeted by crRNAs. After genotyping, expression levels of GADD34 in 2 control and 5 GADD34 KO single cell clones in response to 2 µM thapsigargin (Biotrend) treatment for 3 h was assessed by Western blotting. After characterization and for further experiments, cell pools were generated by mixing the clones in equal proportions.

### Chemical treatments

For UPR^mt^ induction, GTPP (kindly provided by Dr Altieri, The Wistar Institute Cancer Center, Philadelphia USA) was diluted in DMSO and used at concentration of 10 μM for up to 8 h. Paraquat (Sigma-Aldrich, #36541) was dissolved in H_2_O and used at a concentration of 0.5 mM for 48 h. For SG assembly, sodium arsenite (Sigma Aldrich) was diluted in H_2_O and used at a concentration of 0.5 mM for 30 min. Silvestrol (MedChemExpress) was resuspended in DMSO and used at a concentration of 2 μM for 1 h. Phosphorylation of eIF2α was induced by DTT (Promega, DMSO) at 5 mM for 2 h or by UV irradiation at 20 mJ/cm2 using Stratalinker 1800 (Stratagene) followed by 2 h incubation at 37 °C. GCN2 activity was inhibited using 1 μM A-92 (Axon Medchem, dissolved in DMSO). PERK activity was inhibited using 0.1 μM GSK2606414 (Merck, dissolved in DMSO). SG formation was inhibited by pre-incubating cells with G3Ib, or the control drug G3Ib’ (kindly provided by Paul Taylor) at 50 μM final concentration for 20 min before further stimulating with other stressors.

### Transfection with double-stranded (ds) RNA

Synthesis and purification of 200-bp dsRNA was described previously (Klein et al., 2022). In brief, a 200-bp region from the ampicillin resistance gene was used as template for *in vitro* synthesis of plus and minus strand transcripts, hybridized and purified. 2 x 10^5^ cells were seeded in a 6-well plate containing coverslips and transfected with 1 µg dsRNA mixed with Lipofectamine 2000 transfection reagent (Thermo Fisher Scientific) in a 1:3 ratio. Cells were fixed after 16 h with PBS-PFA 4% and further processed for immunofluorescence analysis.

### Immunofluorescence microscopy

Cells were seeded on sterilized coverslips in a 24-well plate and treated as required. Media was removed and fixed with 4% paraformaldehyde (PFA) in PBS for 20 min at RT. Fixation solution was removed, coverslips were rinsed with PB and stored at 4 °C or processed immediately. Cells were permeabilized with 0.1% Triton X-100 (Sigma) in PBS for 5 min at RT and blocked with 0.5% bovine serum albumin (Fisher) in PBS for 1 h at RT. Cells were then incubated with primary antibody solution for 1 h at RT, washed 3 times with PBS and incubated with secondary antibody solution containing 0.2 μg/ml DAPI for 1 h at RT. Coverslips were then rinsed 3 times with PBS and mounted onto microscope slides with 5 μl Mowiol 4-88 (Sigma-Aldrich #81381). Cells were visualized and imaged with a Nikon Ti-Eclipse A1M Confocal Microscope using a 60X objective. Confocal images were analysed utilizing Image J. SG quantification was either performed manually counting at least 100 cells per condition or automatically using the Analyze Particles function, obtaining further information such as the average size of SG and number per cell. Quantification of mitochondria morphology was performed by manually counting cells with distinct mitochondria phenotypes – tubular or fragmented – based on TOMM20 staining. The primary antibodies used were rabbit anti-G3BP1 (1:600, Sigma), goat anti-eIF3η (1:600, Santa Cruz), rabbit anti-TOMM20 (1:1000, abcam ab78547), mouse IgG2a anti-dsRNA (1:250, SCICOBS), mouse IgG1 anti-TOMM20 (1:100, Becton Dickinson), mouse anti-puromycin (1:5, DSHB #PMY-2A4). Secondary antibodies used were goat anti-rabbit Alexa 488, donkey anti-goat Alexa 555, chicken anti-mouse 633, goat anti-mouse-IgG2 Alexa 568 and goat anti-mouse-IgG1 Alexa 488 (1:500, Invitrogen).

### Ribopuromycylation assay

Cells were seeded on sterilized coverslips in a 24-well plate and treated as required. Prior fixation cells were incubated with puromycin (Sigma) at 10 µg/ml at 37° for 5 min to label the nascent polypeptide chains followed by a 2 min pulse at RT with emetine (Sigma) at 180 µM to block translation elongation. Media was removed, cells were rinsed with warm DMEM, fixed with 4% PFA in PBS for 20 min at RT and processed for immunofluorescence as described above. Puromycin incorporation was quantified using ImageJ. The raw integrated density values were measured from ∼ 100 cells per condition.

### Immunoblotting

Cells were seeded in 6-well plates to be around 80-90% confluent at the indicated times. Cells were treated as required, then media was removed, and cells were washed twice with cold PBS. Cells were lysed with protein lysis buffer (50 mM Tris-HCl, 250 mM NaCl, 5 mM EDTA, 1 mM Mg_2_Cl, 1% NP40 and supplemented with protease inhibitor cocktail, Roche), incubated on ice for 20 minutes and spun at 10,000 g for 10 min at 4 °C. Supernatant was collected and protein concentration was measured following BCA pierce kit (Fisher) and between 20-60 μg of protein were used depending on the experiment and quantity obtained. Lysed were mixed with 3x red Loading buffer supplemented with DTT (Cell Signalling) and boiled at 95 °C for 5 min. Lysates were separated in 4-20% gradient gel with 1x SDS running buffer and proteins were transferred to a nitrocellulose membrane using the Pierce Power Station and 1-step Transfer Buffer (Thermo Scientific). Membranes were blocked in 5% BSA or 5% marvel in TBS-Tween (TBS-T) for 1 h at RT and then incubated with primary antibodies overnight at 4 °C. Next day, membranes were washed 3 times with TBS-T and subsequently incubated with secondary antibodies (Dako, 1:5000) diluted in 5% marvel in TBS-T for 1 h at RT. After three washes with TBS-T, secondary antibodies were labelled with Clarity Western ECL Substrate (Bio-Rad), and membranes were imaged using the vilber-Lourmat Imaging system. Band intensity was quantified and analysed using Image Lab Software (Bio-rad). Primary antibodies used were rabbit anti-eIF2α (1:1000, Cell Signaling), rabbit anti-P-eIF2α (1:1000, Invitrogen), rabbit anti-ATF4 (1:1000, Cell Signaling), rabbit anti-GADD34 (1:1000, Proteintech), mouse anti-GAPDH (1:5000, Ambion), goat anti-eIF3η (1:2000, Santa Cruz) and anti-eIF3A (1:2000, Bethyl Laboratories).

### Preparation of RNA samples and RT-qPCR

Cells were seeded in 6-well plates to be around 80-90% confluent at the indicated times. Cells were treated as needed, then were washed twice with cold PBS, and total RNA was extracted using a quick-RNA micro prep kit (Zymo Research) following the manufacturer’s instructions. Purified RNA was quantified on Nanodrop, and 500 ng of RNA was subject to reverse transcription using Precision nanoScript2 Reverse Transcription kit from Primer Design. Next, real-time quantitative PCR (qPCR) was performed in duplicate with 25 ng of template, specific primers listed below used at a final concentration of 0.3 μM and Precision Plus Master Mix (Primer Design) following manufacturer’s instructions, using the Quant Studio 7 Flex (Applied Biosystems). The comparative Ct method was used; Ct values from genes of interest were normalized to RpL9 Ct values unless otherwise specified. The 2-ΔΔCT method was used to calculate the fold change expression relative to non-treated (Livak and Schmittgen, 2001).

### Mitochondrial respiration

Oxygen consumption rate of U2OS WT and G3BP1/2 dKO cells was measured on an extracellular flux analyzer (Agilent Seahorse) using the XF Cell Mito Stress Test Kit following manufacturer’s protocol. Briefly, cells were seeded one day prior to the experiment on Seahorse XFe24 culture plate at 80% confluency. Cells were treated as required and 1 h prior running the plate, medium was removed, and cells were incubated with assay medium supplemented with 1 mM pyruvate, 2 mM glutamine and 10 mM glucose. Cells were then sequentially stimulated with oligomycin at 1 µM, FCCP at 2 µM and rotenone and antimycin A at 1 µM and respiratory profile was obtained. Data was normalised by amount of protein for each well. Following assay, cells were lysed with protein lysis buffer (50 mM Tris-HCl, 250 mM NaCl, 5 mM EDTA, 1 mM Mg_2_Cl, 1% NP40 and supplemented with protease inhibitor cocktail, Roche), incubated on ice for 20 minutes and spun at 10,000 g for 10 min at 4 °C. Supernatant was collected and protein concentration was measured following BCA pierce kit (Fisher).

### Mitochondrial ROS measurement

Cells were seeded in 96-well plate to be around 60% confluent for the day of the experiment. After GTPP treatment for 2 h, cells were washed and incubated with 2.5 μM MitoSOX Red (Invitrogen, #M36008) diluted in warm PBS for 30 min at 37 °C. Next, cells were washed three times and fluorescence was analysed in IncuCyte S3 imaging system. MitoSOX Red mean intensity was extracted from 4 images taken at 20X per condition and plotted relative to DMSO-treated cells.

### Mitochondrial translation assay

To label newly synthesized mitochondrial-encoded proteins, a click chemistry-based approach was applied (Yousefi et al., 2021). Briefly, WT and G3BP1/2 dKO U2OS cells were treated with 10µM GTPP or DMSO for 2 h, before replacing existing media with methionine-free media (D0422, Sigma) supplemented with 2nM L-glutamine. Cytoplasmic translation was inhibited with cycloheximide (50µg/ml, 20 mins), before addition of methionine analogue L-Homopropargylglycine (HPG) which is preferentially incorporated into nascent mitochondrial peptides, for 15 mins. To validate this approach, cells were left in methionine-containing media, whereby methionine was preferentially incorporated into mitochondrial ribosomes instead of HPG, resulting in loss of HPG fluorescence signal. For control samples, mitochondrial translation was blocked using Chloramphenicol (CHL, 150 µg/ml, 50 mins) before HPG labelling. Cells were then incubated in buffer A (10 mM HEPES, 10 mM NaCl, 5 mM KCl, 300 mM sucrose, and 0.015% digitonin) for 2 min on ice, followed by 15 s in buffer A without digitonin before fixation with 4% PFA in PBS for 30 min at RT. Next, click chemistry and immunofluorescence was performed. After fixation, coverslips were rinsed in PBS for 5 min then quenched with 100 mM NH4Cl in PBS for 15 min and blocked with 5% BSA in PBS for 10 min. Click chemistry was performed as per the manufacturer guidelines. Briefly, coverslips were then incubated with 500µl Picolyl azide AF-555 (CLK-090-1, Jena Bioscience) in CuAAC BTTAA Cell Reaction Buffer (CLK-073, Jena Bioscience) in the dark for 60 min at RT. Coverslips were washed twice with 3% BSA in PBS for 5 min before IF staining. The probes used in this experiment were TOMM20 (AB78547, Abcam) and chicken anti-mouse Alexa fluor 488 (A21200, life technologies).

### Cell viability assay

Cells were seeded in 96-well plate, incubated with GTPP and Cytotox dye (Sartorius, #4632) at a final concentration of 250 nM and monitored overtime using IncuCyte S3 live cell analysis system. When plasma membrane integrity is disrupting in dying cells, IncuCyte Cytotox Dye enters cells and massively increases its fluorescence upon DNA binding. About 9 pictures were taken at 10X per condition in 96-wells every 2-3 h over a 28 h period.

### Statistical analyses

Statistical analyses were performed using Graphpad Prism software and three biological replicates were analysed in all experiments. Statistical tests performed are indicated in the figure legends (****, P < 0.0001; ***, P < 0.001; **, P < 0.01; *, P < 0.05; ns, not significant).

### Primer list

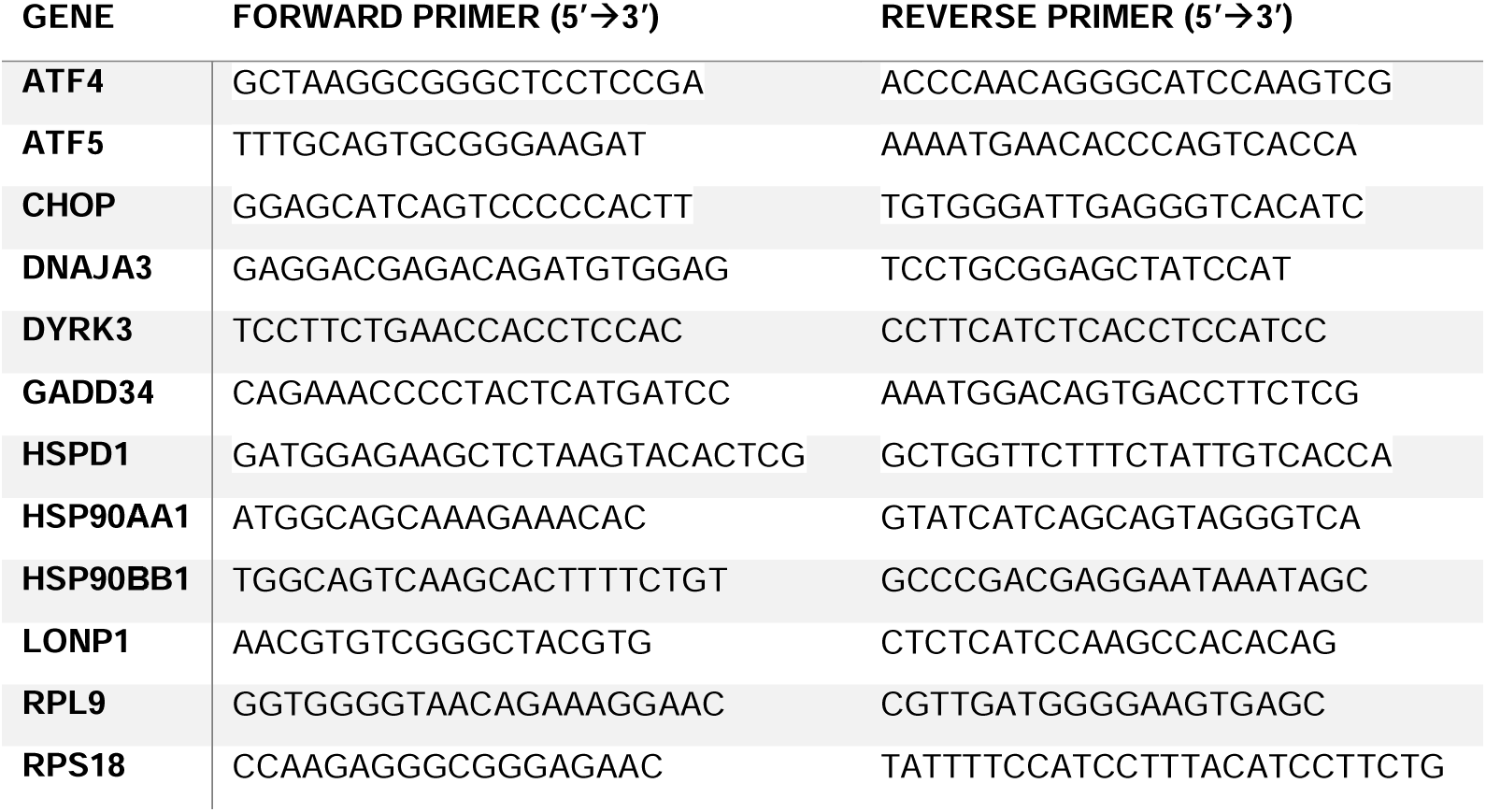
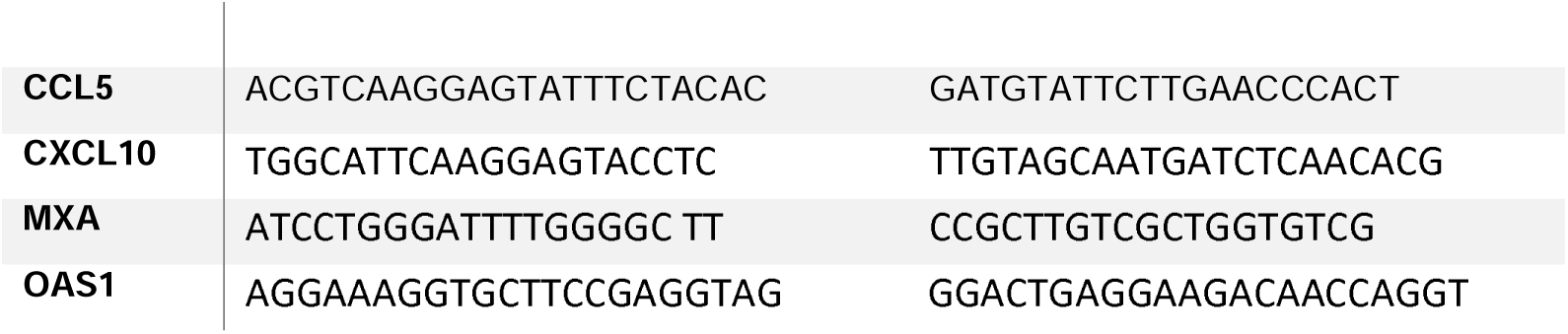

## Figure Legends

**Supplementary Figure 1: Inhibition of SGs formation by G3Ib blocks GTPP-induced SG assembly.** U2OS cells were preincubated with either DMSO or G3Ib/G3Ib’ at 50 μM for 20 min and GTPP was added to the cells at final concentration of 10 μM. SG formation was analysed by immunofluorescence using SG marker G3BP1 (cyan). Nuclei were stained with DAPI. Scale bars indicate 10 μm.

**Supplementary Figure 2: Translational shut off upon UPR^mt^ activation by GTPP.** U2OS WT cells were stimulated with GTPP for the indicated times or arsenite for 1 h and treated with puromycin followed by emetine before fixation. Then, cells were analysed by immunofluorescence with anti-puromycin (grey) and the SG markers anti-G3BP1 (cyan) and anti-eIF3η (magenta). Nuclei were stained with DAPI. Scale bars indicate 10 μm.

**Supplementary Figure 3: GTPP treatment does not result in mitochondrial dsRNA leakage.** Cells were treated with GTPP for 2 h or transfected with dsRNA (positive control) and analysed by immunofluorescence with anti-dsRNA (magenta) and the mitochondrial marker anti-TOMM20 (cyan). Nuclei were stained with DAPI. Scale bars indicate 10 μm.

**Supplementary Figure 4: GTPP treatment impairs subsequent SG formation by arsenite.** Individual panels are shown from Fig 3A.

**Supplementary Figure 5: Paraquat treatment results in UPR^mt^ activation but does not induce SG formation in U2OS cells. (A)** RT-qPCR analysis of ISR and UPR^mt^ markers in U2OS cells treated with 0.5 mM paraquat for 48 h. Results shown as mean ± SD, n=3, normalised to RPL9 mRNA and shown relative to NT expression level, analysed by two-way ANOVA (****, P < 0.0001; *, P < 0.05); NT, non-treated. (**B**) Treatment of U2OS cells with paraquat for 1, 2, 6, 24 and 48 h does not result in SG assembly. Representative images from three independent experiments are shown. Cells were analysed by immunofluorescence for the SG markers G3BP1 (cyan) and eIF3η (magenta); nuclei were stained with DAPI. Scale bars indicate 10 μm. (**C**) Individual panels are shown from Fig 3E.

**Supplementary Figure 6: ATF5 silencing modulates SG dynamics in a stress-specific manner.** (**A**) U2OS cells were transfected with non-targeting siRNA (siCont) or ATF5-specifc siRNA (siATF5), after 48 h cells were treated with GTPP 4 h and sodium arsenite (Ars) was added 30 min before harvesting and cells were analysed by immunofluorescence for the SG markers G3BP1 (cyan) and eIF3η (magenta). Nuclei were stained with DAPI. Scale bars indicate 10 μm. (**B**) Panel shows quantification by manual counting of cells with SGs using ImageJ, at least 100 cells were counted per replicate. Results are shown as mean ± SD, n=3 and were analysed by two-way ANOVA (ns, non-significant). (**C, D**) Violin plot displaying the average size of SGs per cell and the number of SGs per cell from GTPP treated cells (**C**) and GTPP+Ars treated cells (**D**) quantification was performed using G3BP1 channel and Analyze Particle plugin from ImageJ. Figure shows the median and quartiles of at least 60 cells per condition from three independent experiments and were analysed by Mann-Whitney test (**, P < 0.01).

**Supplementary Figure 7: Validation of U2OS GADD34 KO single cell clones by Western blot. (A)** Clones were treated with 2 µM thapsigargin (Thap) or with DMSO (-) for 3 h and GADD34 protein levels were analysed by western blot. eIF3A was used as loading control. (**B, C**) Individual panels are shown from Fig 4A and 4C, respectively.

**Supplementary Figure 8: U2OS G3BP1/2 dKO cells do not form SGs.** U2OS WT and G3BP1/2 dKO cells were treated with arsenite (Ars) for 30 min or GTPP for 2 h and analysed by immunofluorescence using the SG markers G3BP1 (cyan) and eIF3η (magenta). Nuclei were stained with DAPI. Scale bars indicate 10 μm.

**Supplementary Figure 9: U2OS G3BP1/2 dKO cells presented reduced MitoSOX Red signal after GTPP treatment compared to WT cells.** Representative IncuCyte images from Figure 6C showing MitoSOX Red and phase channels.

## Data availability Statement

All the data are included in the article. All unique reagents generated in this study are available from the corresponding authors with a completed Materials Transfer Agreement.

## Acknowledgments

We thank P. J. McCormick (Queen Mary University London) for helpful discussions during the study.

## Author contributions

N. L., A. R. and I. S. conceptualization; N. L., M. L-N., A. R., B.D.F., J.P.T., Z. S., R. S. and I. S. methodology; N. L., M. L-N., A. R., and Z. S. formal analysis; N. L., M. L-N., E.R., A. R., Z. S., R. S. and I. S. investigation; N. L. and M. L-N. writing–original draft; N. L., M. L-N., A. R., E.R., Z. S., R. S., B.D.F., J.P.T. and I. S. writing–review & editing; N.L. supervision. N. L., A. R., and I. S. funding acquisition.

## Funding and additional information

Work in N.L.’s laboratory is supported by the Biotechnology and Biological Sciences Research Council research and strategic grants BB/V014528/1 and BBS/E/PI/230002A. Bioimaging studies are supported by BBSRC Core Capability Grant BBS/E/I/00007039. M. L-N. and R. S. are supported by DCSA4 Ph.D studentships from the University of Surrey. Work in A.R.’s laboratory was supported by the Deutsche Forschungsgemeinschaft (DFG, German Research Foundation) project numbers 240245660 SFB1129 TP13 and 278001972 TRR186 A14.

