## Supplementary figures and images for "Activation of the mitochondrial unfolded protein response regulates the formation of stress granules"

### supplementary figure 1

**Supplementary Figure 1**

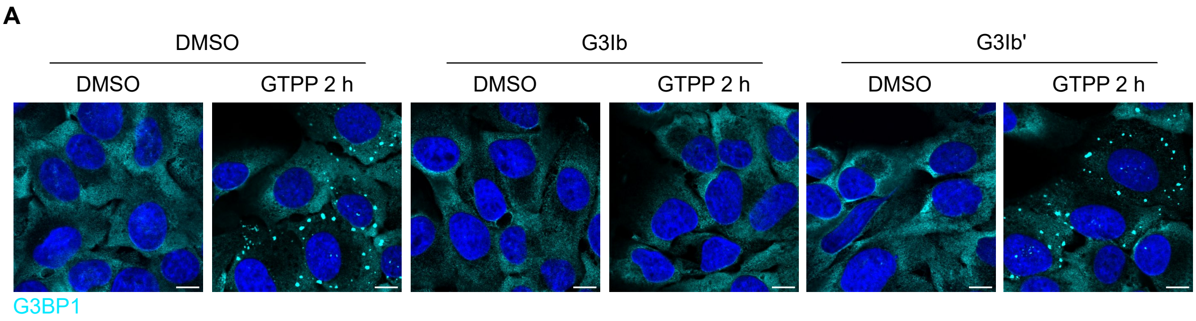

### supplementary figure 2

**Supplementary Figure 2**

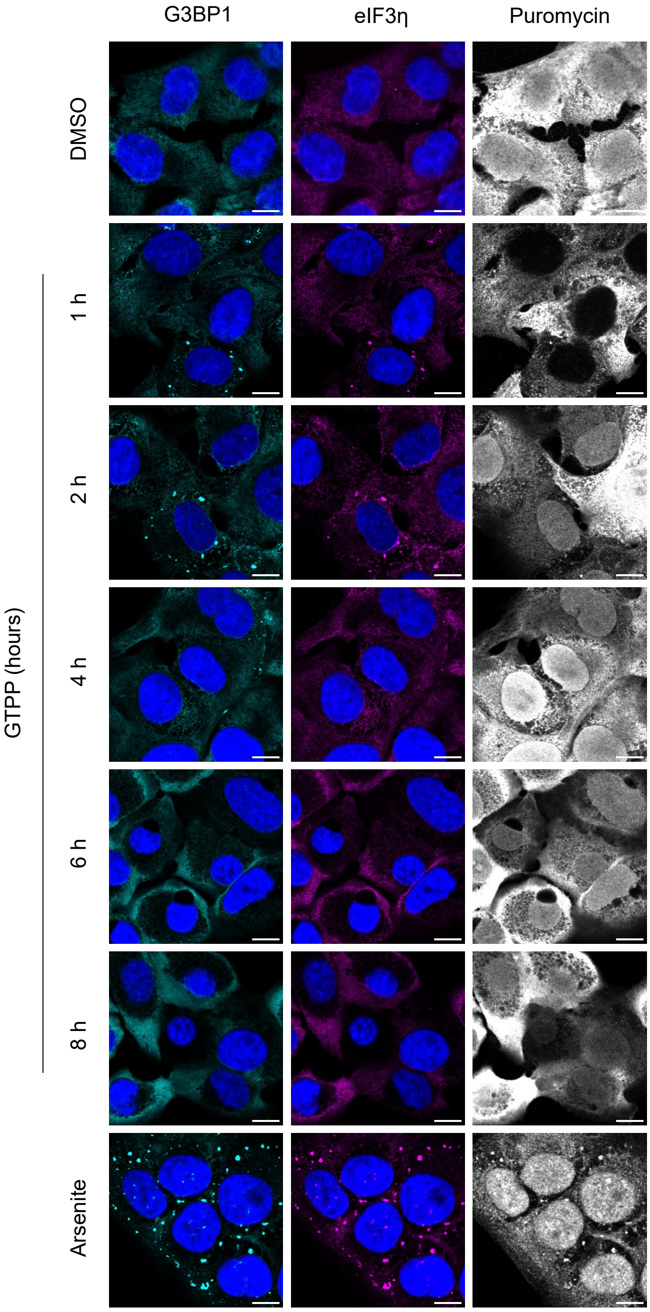

### supplementary figure 3

**Supplementary Figure 3**

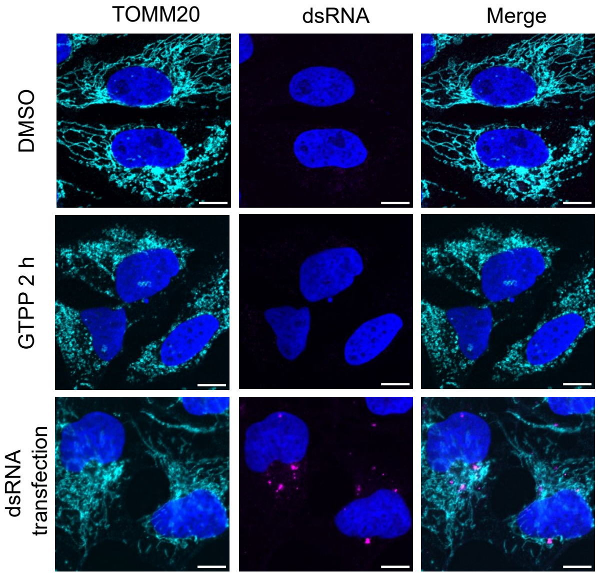

### supplementary figure 4

**Supplementary Figure 4**

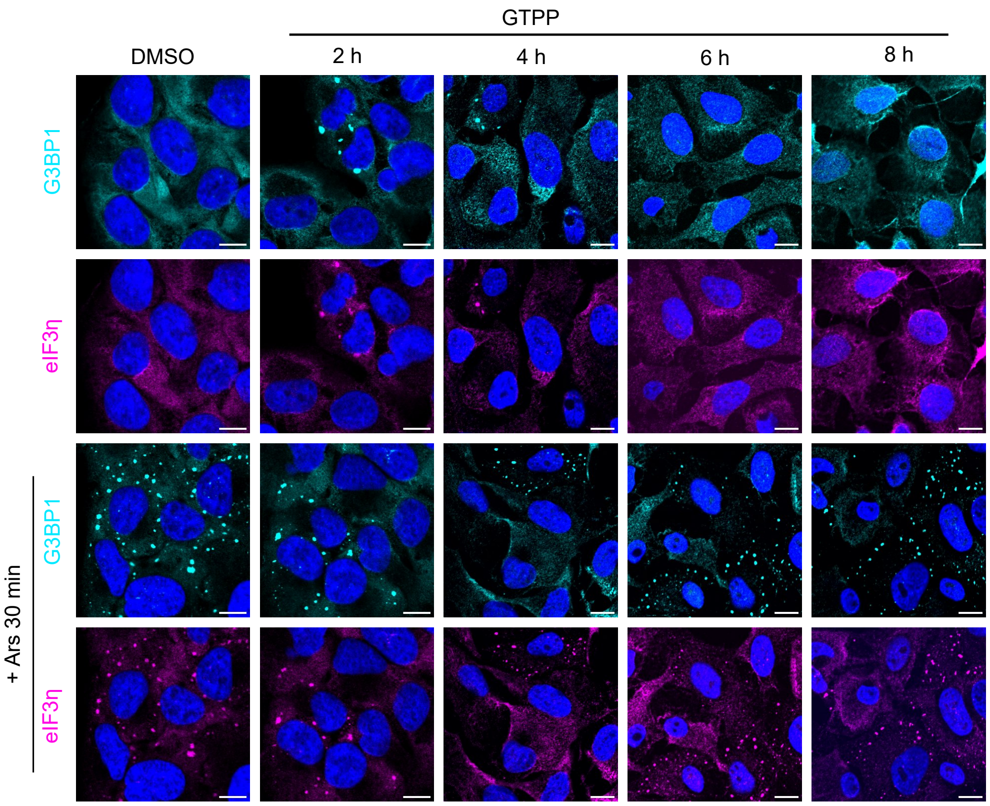

### supplementary figure 5

**Supplementary Figure 5**

A

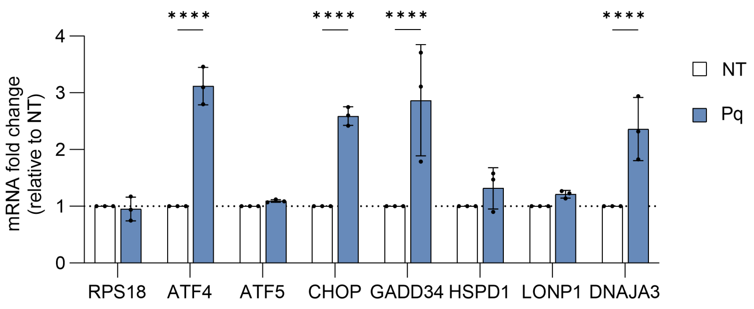


B

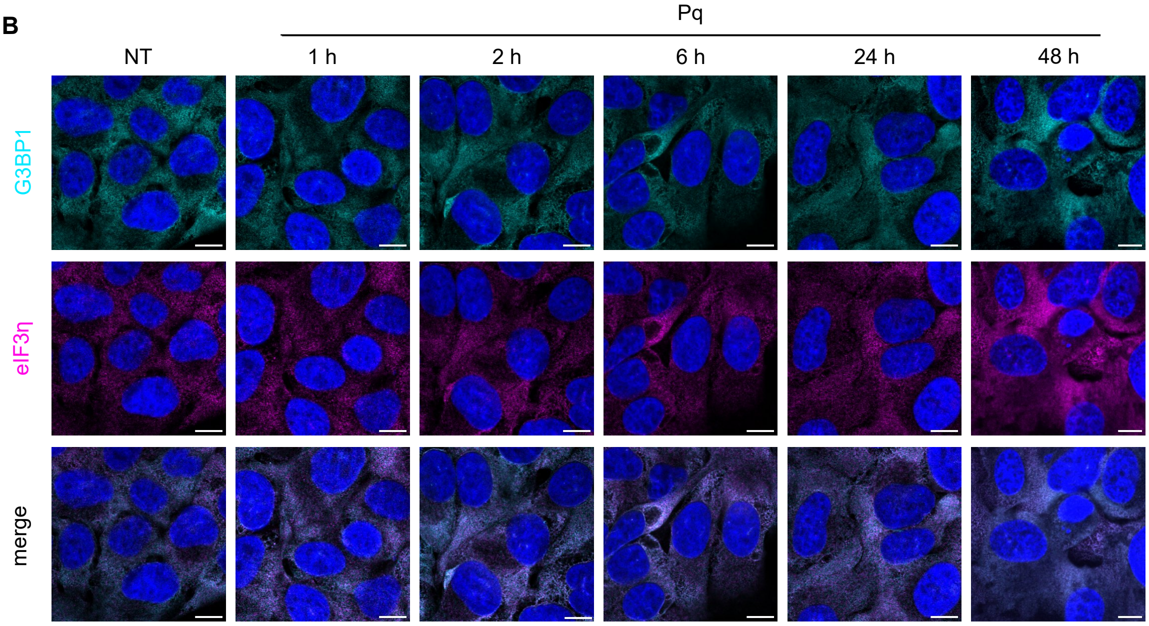


C

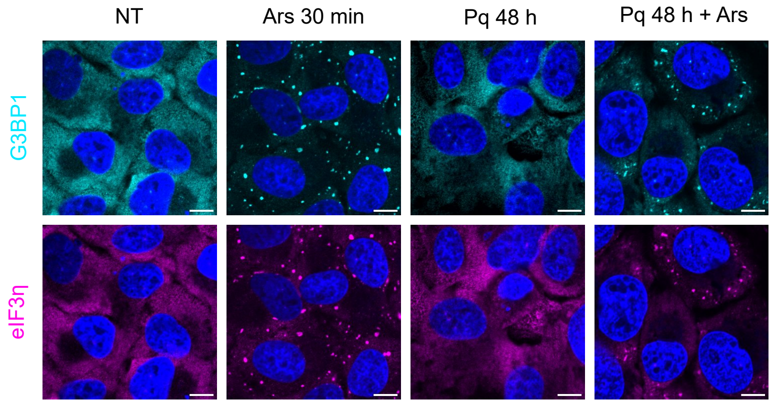

### supplementary figure 6

**Supplementary Figure 6**

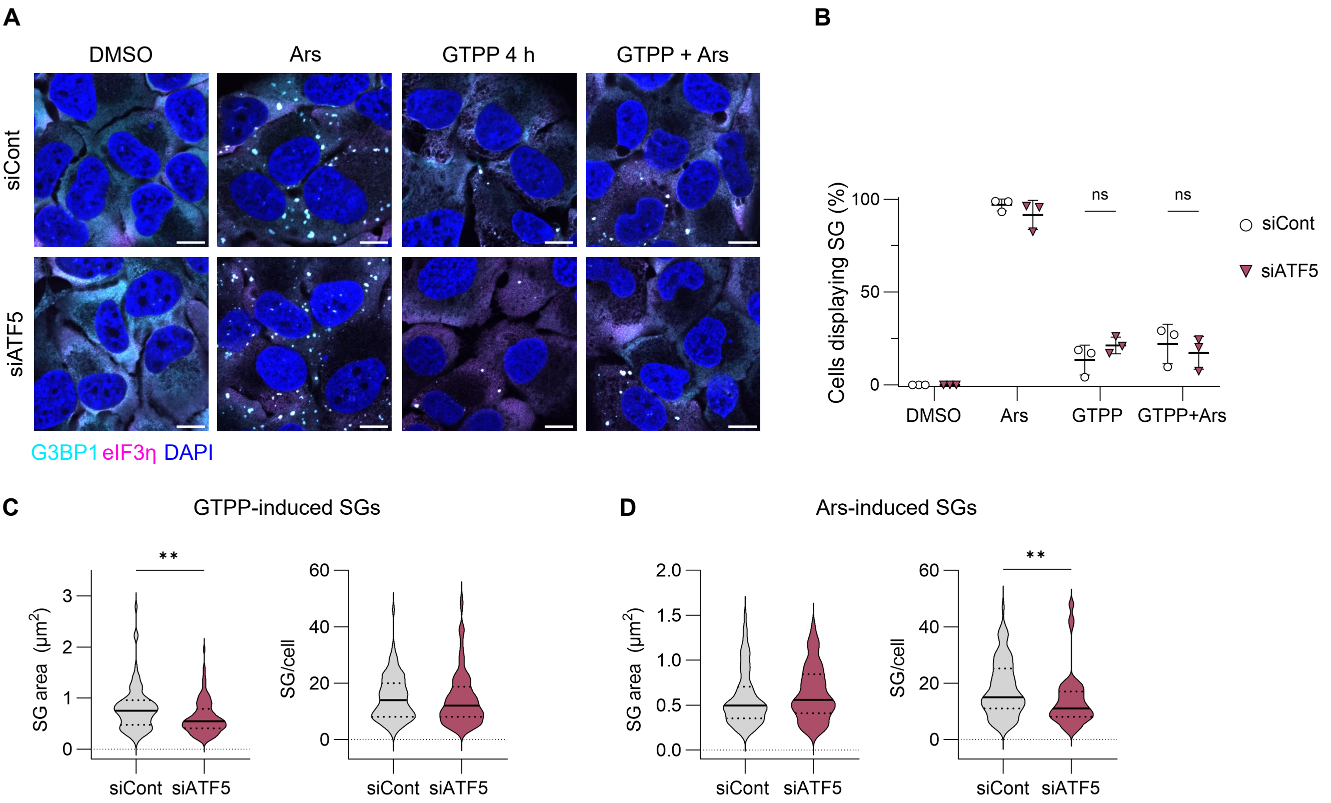

### supplementary figure 7

#### Supplementary Figure 7

**A**

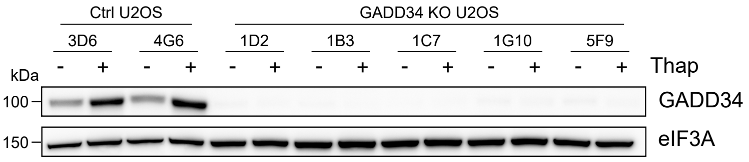


**B**

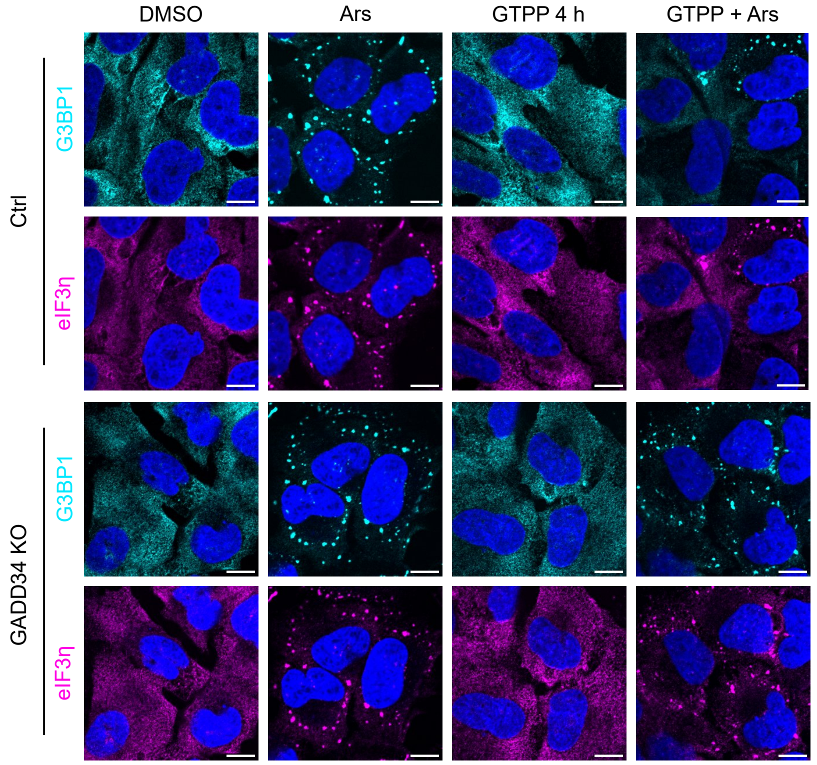


**C**

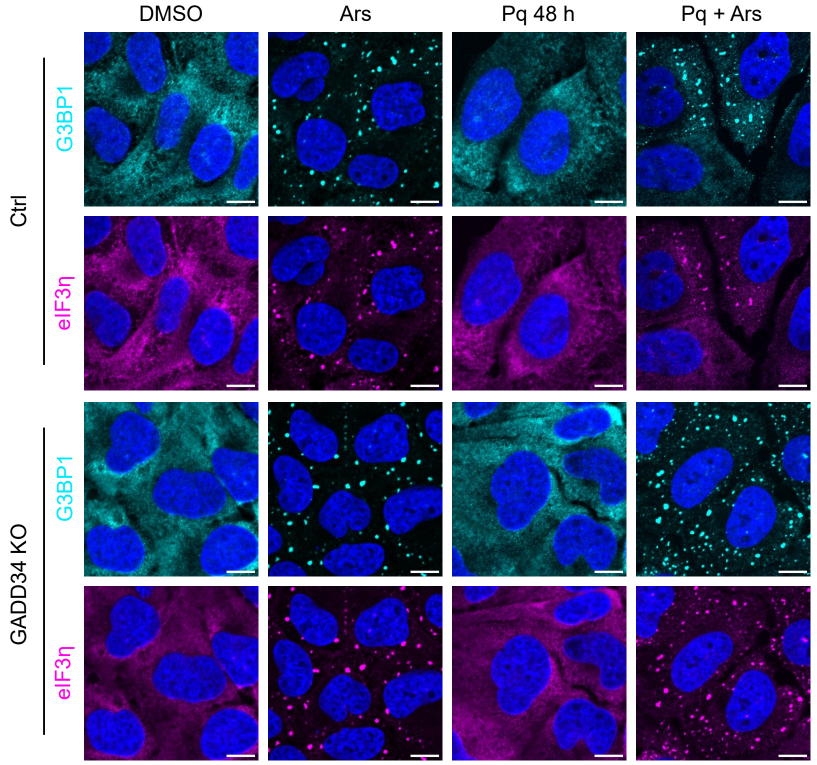

### supplementary figure 8

**Supplementary Figure 8**

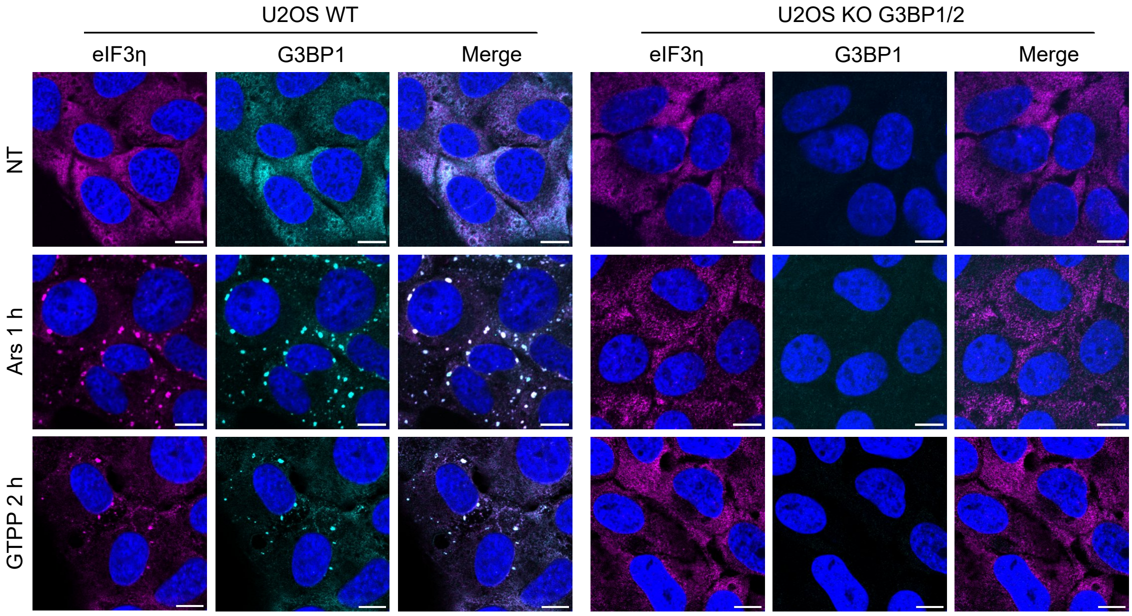

### supplementary figure 9

**Supplementary Figure 7**

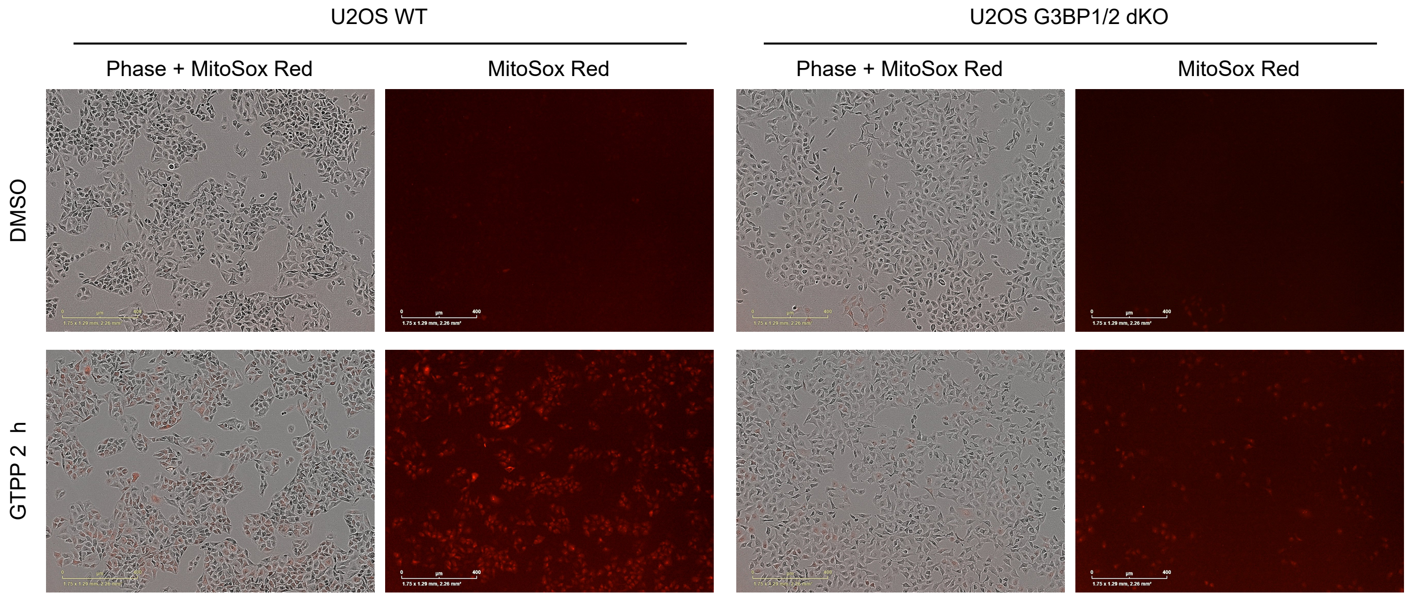
